# Bacterial hemophilin homologs and their specific type eleven secretor proteins have conserved roles in heme capture and are diversifying as a family

**DOI:** 10.1101/2023.03.15.532718

**Authors:** Alex S. Grossman, David A. Gell, Derek G. Wu, Heidi Goodrich-Blair

## Abstract

Cellular life relies on enzymes that require metal cofactors, which must be acquired from extracellular sources. Bacteria utilize surface and secreted proteins to acquire such valuable nutrients from their environment. These include the cargo proteins of the type eleven secretion system (T11SS), which have been connected to host specificity, metal homeostasis, and nutritional immunity evasion. This Sec-dependent, Gram-negative secretion system is encoded by organisms throughout the phylum Proteobacteria, including human pathogens *Neisseria meningitidis, Proteus mirabilis, Acinetobacter baumannii,* and *Haemophilus influenzae*. Experimentally verified T11SS-dependent cargo include host metal acquisition proteins <u>t</u>ransferrin <u>b</u>inding <u>p</u>rotein B (TbpB) and lactoferrin <u>b</u>inding <u>p</u>rotein B (LbpB), as well as the hemophilin homologs <u>h</u>eme <u>r</u>eceptor <u>p</u>rotein C (HrpC) and <u>h</u>emo<u>ph</u>ilin A (HphA), the complement immune evasion protein factor-<u>H b</u>inding <u>p</u>rotein (fHbp), and the host symbiosis factor <u>n</u>ematode intestinal localization protein C (NilC). Secretion of each of these cargo proteins relies on a specific T11SS. Here, we examined the specificity of T11SS systems for their cognate cargo proteins using taxonomically distributed homolog pairs of T11SS and hemophilin cargo and explore the ligand binding ability of those hemophilin homologs. Our comparative ligand binding analysis of four hemophilin family proteins identified previously unknown ligand binding diversity within this protein family, which informed our description of structural features that are likely to contribute to heme/porphyrin binding specificity. *In vivo* expression of hemophilin homologs revealed that each was secreted in a specific manner by its cognate T11SS protein. Furthermore, secretion assays of chimeric hemophilin proteins revealed that specificity is predominantly dictated by the C-terminal domain of the cognate cargo. Meanwhile, the N-terminal effector domains of these T11SS-dependent cargo proteins feature porphyrin binding pockets that drive ligand binding affinity and specificity. In light of these results, we have termed this N-terminal domain the hemophilin ligand binding domain (Hlb) after its first characterized representative.

## Introduction

Many enzymes have evolved expanded catalytic potential through the incorporation of metallic cofactors and prosthetic groups. Iron cofactors are essential to most living organisms due to their functional contributions to enzymes required for DNA synthesis, photosynthesis, respiration, and nitrogen metabolism [1]. Without the biochemical flexibility provided by metallic cofactors, life as we know it would be impossible. Because of this, and the limited bioavailability of essential metals, competition among organisms for these ions can be fierce. In some cases, including many marine environments, competition for bioavailable iron is actually the major limiting factor of microbial growth [2]. This phenomenon is exploited by animals through a process known as nutritional immunity, in which valuable ions, such as iron, are sequestered to slow or deter the pathogenic growth of microbes. Within animal hosts iron is sequestered by proteins such as hemopexin, transferrin, lactoferrin, and ferritin [3]. Medical conditions, such as hemochromatosis, that increase the serum iron concentration or prevent effective storage of iron increase a patient’s risk of infection from many bacteria, including *Escherichia coli*, *Listeria monocytogenes*, and *Yersinia enterocolitica* [4]. In turn, to overcome iron limitation within an animal host, bacteria have evolved means of countering nutritional immunity, including adaptations to use alternative catalytic metals [5], production of high-affinity siderophores [6], and or membrane bound uptake receptors [3] that facilitate acquisition of iron from host metalloproteins.

Several recent studies have linked the <u>t</u>ype <u>eleven s</u>ecretion <u>s</u>ystem (T11SS) <u>o</u>uter <u>m</u>embrane <u>p</u>roteins (OMPs) and their cargo proteins to iron uptake strategies in Gram-negative bacteria. In *Neisseria*, the T11SS proteins Slam1 and Slam2 surface expose cargo proteins that are responsible for binding host-metal carriers: Transferrin binding protein B (TbpB) and Lactoferrin binding protein B (LbpB) are surface exposed by Slam1, and Hemoglobin/Haptoglobin binding protein A (HpuA) is surface exposed by Slam2 [7–10]. These surface exposed outer membrane lipoproteins facilitate bacterial colonization by capturing their respective host factors (transferrin, lactoferrin, or hemoglobin/haptoglobin) and complexing with a TonB-dependent uptake channel capable of importing the iron cofactor. Since surface exposure is essential for the function of these lipoproteins, genetic inactivation of the T11SS OMP, Slam1 prevents effective colonization and pathogenesis by *Neisseria* [11]. While *Neisseria* Slam1 and Slam2 have specificity for their respective cargo [7], no underlying mechanism for specificity has been proposed yet and it is unknown if all T11SS have specificity for their cognate cargo. Bioinformatic analyses revealed a large number of potential T11SS- dependent cargo, lipid anchored and unanchored, which frequently exist in cognate pairs/groupings according to genomic co-occurrence analyses [11,12]. To date, all verified or predicted T11SS-dependent cargo have two distinct domains: an N-terminal domain that varies in predicted structure and function across cargo, and a C-terminal, 8-stranded β-barrel domain from the TbpBBD or lipoprotein C families, though the latter is not exclusive to lipoproteins but also occurs in non-lipidated proteins [7,11,12].

T11SS are capable of secreting unlipidated cargo proteins, such as the soluble hemophores <u>h</u>eme <u>r</u>eceptor <u>p</u>rotein C from *X. nematophila* (HrpC) and <u>h</u>emo<u>ph</u>ilin A from *A. baumanii* (HphA) [12,13]. HphA likely captures heme from hemoglobin and other host hemoproteins, and contributes to the virulence of *Acinetobacter baumannii* in a murine infection model through its role as a co-receptor to the TonB-dependent heme receptor HphR [13]. Thus, hemophilin proteins represent a high-affinity heme acquisition system comparable to HasA from *Serratia marcescens* [14] or IsdB from *Staphylococcus aureus* [15]. Known members of the hemophilin protein family function to import heme from a host environment as depicted in Figure 1. Compiling the results of published data from multiple organisms into a single model suggests that that hemophilin crosses the inner membrane through the Sec translocon to reach the periplasm [12], may interact with chaperones such as Skp to traverse the periplasm [16], and then crosses the outer membrane in a T11SS-dependent manner to reach the extracellular milieu [7,12,13,17]. From here, hemophilin captures heme that is released from hemoglobin or other host hemoproteins using a high affinity binding domain [13,18]. Holo-hemophilin is predicted to interact with a TonB-dependent outer membrane co-receptor which imports the heme molecule into the periplasm and releases apo-hemophilin. Finally, the hemin utilization system delivers periplasmic heme into the cytoplasm for incorporation into cellular processes or digestion by heme oxygenase to free the iron cation [19].

**Figure 1.**
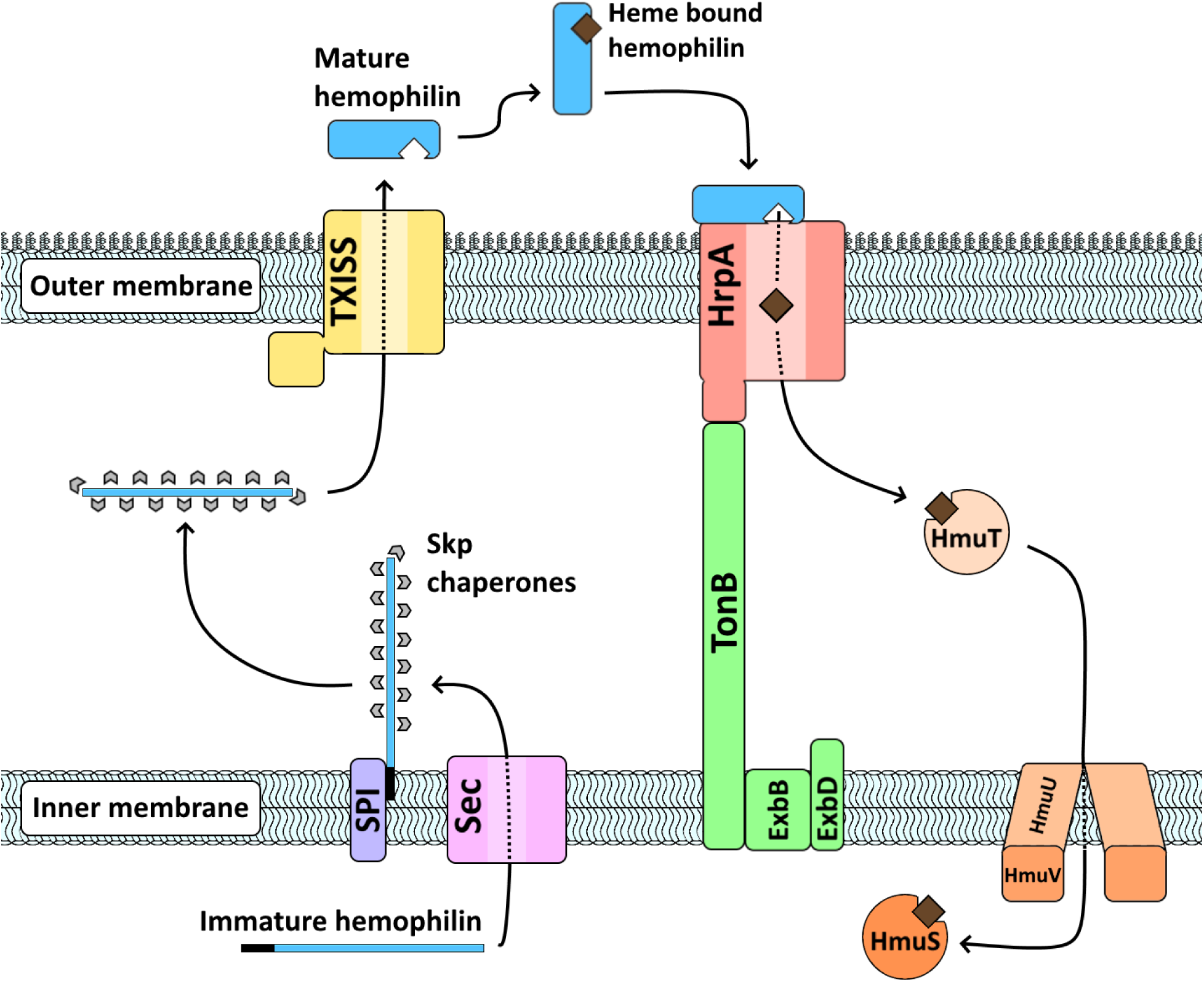
Conceptual model of hemophilin mediated heme acquisition. Described hemophilin homologs have a signal peptide directing them to the Sec translocon and signal peptidase I. Based on analogy to the T11SS-dependent lipoprotein TbpB, hemophilin may depend on Skp for periplasmic transport. Each hemophilin protein has a cognate T11SS which translocates it through the outer membrane where it acts to bring heme to a TonB-dependent receptor for uptake. Based on analogy to the *Yersinia pestis* Has hemophore system, heme is transported through the periplasm and across the inner membrane by the hemin utilization system (Hmu). Once brought into the cytoplasm, heme can be degraded by heme oxygenase (HmuS), an enzyme which is frequently encoded adjacent to hemophilin loci.

Within this overall framework, hemophilin can have diverse functional roles within different organisms and environments. For example, in *Haemophilus haemolyticus* hemophilin can act as a probiotic factor by making bioavailable iron inaccessible to nontypeable *Haemophilus influenzae* [20]. Hemophilin producing strains of *H. haemolyticus* inhibit the growth of *H. influenzae* significantly more than non-hemophilin encoding strains in co-cultured media [18] and cell cultures [20]. Additionally, oropharyngeal sampling of human subjects indicates that individuals who carry hemophilin encoding *H. haemolyticus* are between 2.43 and 2.67 times less likely to carry nontypeable *H. influenzae* [21]. Conversely, within the opportunistic pathogen *A. baumannii,* hemophilin can act as a virulence factor by facilitating systemic infection in a murine model [13]. Furthermore, predicted hemophilin homologs found in sequence databases display sequence variation within the heme-binding handle domain, suggesting possible variability in ligand binding. To better understand the fundamental biochemical and biological functions of hemophilin family proteins we sought to expand our understanding of hemophilin secretion and activities across a broader taxonomic range.

## Results

### Sequence similarity networks reveal hemophilin families that have genomic associations with metal-related metabolic functions

To explore the relatedness of hemophilin family proteins and to identify subcluster divisions that may reflect divergent function, a sequence similarity network generated through Enzyme Function Initiative-Enzyme Similarity Tool (EFI-EST) was overlaid with a taxonomic framework [22] (Fig. S1 and Supplemental File 1). The network was populated with the previously published dataset of T11SS-associated cargo that were not predicted to be lipidated or membrane anchored [12]. This analysis revealed a single major cluster containing all previously described hemophilin proteins (88/107 nodes), one smaller cluster containing uncharacterized proteins from predominantly *Pseudomonas* and *Neisseria* species (9/107 nodes), and a few unassociated doublets and singletons (10/107 nodes). To focus this study specifically on hemophilin and its direct evolutionary relatives, all nodes not within the central cluster were removed. The remaining nodes were labeled according to taxonomic family and separated using a force-directed separation algorithm, resulting in five subclusters predominantly populated by seven families from Alphaproteobacteria, Betaproteobacteria, and Gammaproteobacteria (Fig. 2AC). The three subclusters that contained Hpl, HrpC, and HphA, respectively were named Hpl-like, HrpC-like, and HphA-like after their respective characterized member. One novel subcluster was termed ‘Cobalt/molybdenum associated’ due to its genomic co-occurrence with genes predicted to encode cobalt- or molybdenum-dependent enzymes. Another novel subcluster was termed ‘Plant/Environmental’ due to the dominant presence of homologs encoded by microbes found in soil, water, and plant-associated environments. The subclusters did not fall exclusively along taxonomic lines. For instance, the Hpl-like subcluster includes *Neisseria* and *Pasteurella* hemophilin homologs that clustered closely together, possibly indicating that these genes may have been horizontally exchanged. Cross-taxonomic-class sequence similarity also was observed in the Plant/Environmental cluster of hemophilin homologs.

**Figure 2.**
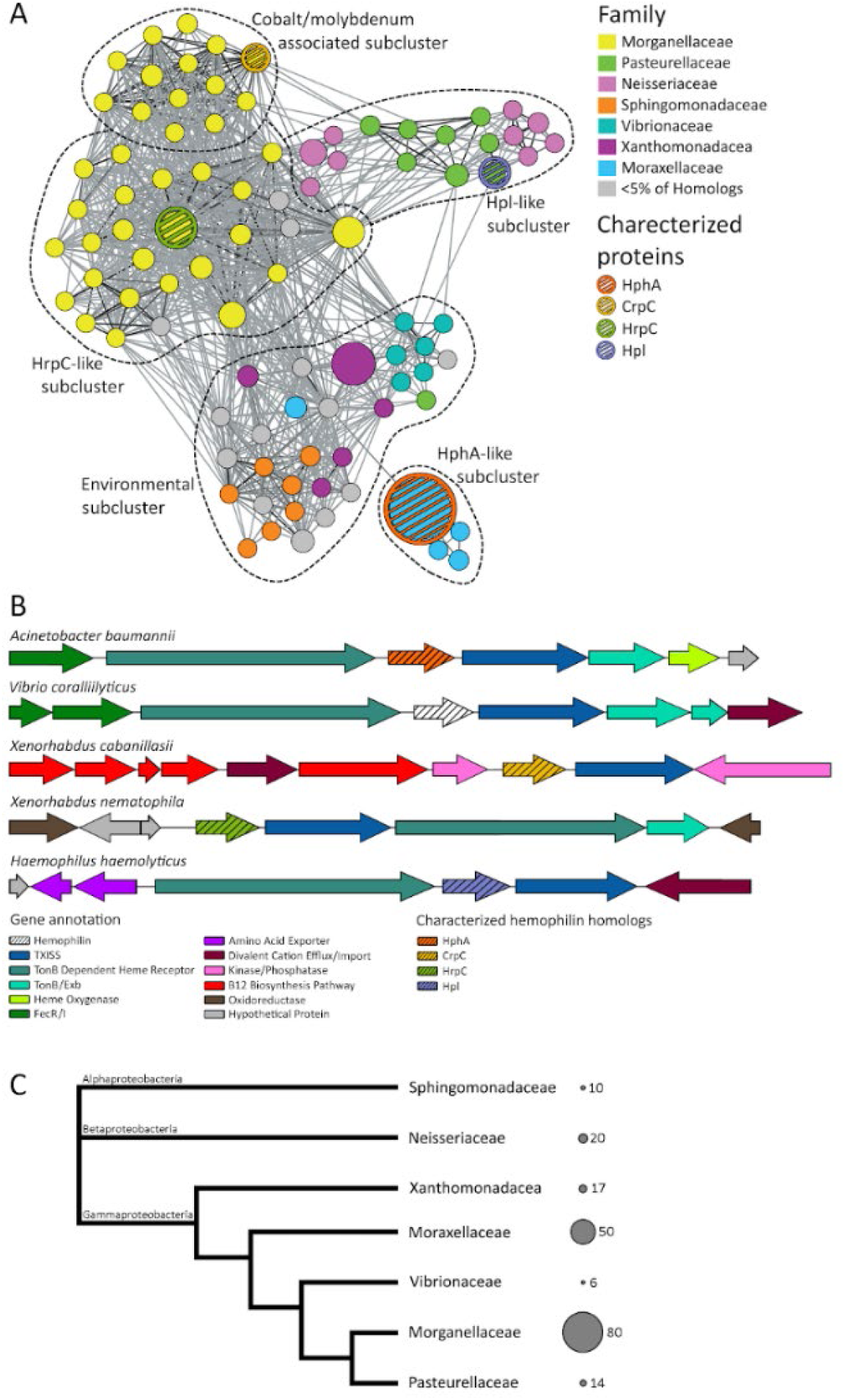
Distribution and relatedness of hemophilin family proteins. **A)** A sequence similarity network of hemophilin homologs generated with EFI-EST. Each node represents one or more protein sequences with 80% or greater identity, the larger the node the more sequences it contains. Edges indicate an alignment score of 35 or greater. Edge darkness indicates shared sequence identity, with the darkest edges being the most identical. Dotted lines indicate proposed subclusters as defined using a force-directed algorithm and the distribution of characterized proteins. **B)** Representative genomic neighborhoods from the subclusters identified within the sequence similarity network demonstrating common co-occurring genes. **C)** A cladogram of the seven genera which encode the most hemophilin family proteins, split between Alpha-, Beta-, and Gammaproteobacteria. The circles and numbers to the right of the cladogram indicate the relative abundance of known hemophilin homologs encoded by each genus.

To identify potential distinguishing features among the five subclusters, we examined the identities of genes that commonly occur within genomic neighborhoods (± 6 genes) with *hrpB*. All sequences from the network were submitted to the Rapid ORF Detection & Evaluation Online (RODEO) web tool that uses profile hidden Markov models to identify co-occurring protein domains (Table S1) [23]. Consistent with previous observations, 92.3% (240/260 sequences) of hemophilin homologs, regardless of subcluster, were encoded in association with genes predicted to encode TonB and TonB-dependent receptors, suggesting a strong and consistent link between T11SS cargo and TonB-dependent uptake across the outer membrane [11,12]. However, some subclusters had further informative co-occurrences. For example, hemophilin homologs in the HphA-like subcluster showed nearly universal co-occurrence with genes predicted to encode heme oxygenase (51/53) and the iron-sensing regulator FecR (48/53). Similarly, homologs in the Plant/Environmental subcluster typically were encoded near genes predicted to encode metal responsive regulatory proteins (FecR or Fur) (64/72) and occasionally near genes encoding heme oxygenase (19/72). Additionally, 69/72 Plant/Environmental-subcluster loci encoded additional regulatory genes such as RpoE, IscR- family regulators, and LysR-family regulators. RpoE is a sigma factor that responds to extra- cytoplasmic stress and is essential for metal resistance in *E. coli* [24]. IscR regulates iron-sulfur cluster biosynthesis according to cellular demand [25], and LysR-family regulators drive diverse pathways by binding DNA directly in response to co-inducing/co-repressing ligands [26]. Homologs in the Cobalt/molybdenum-associated subcluster occurred alongside other predicted T11SS-dependent cargo (14/18) and were either located adjacent to a B12 biosynthetic locus (3/18) or near a formate dehydrogenase locus (13/18). Formate dehydrogenase activity relies on molybdenum metal cofactors, and their co-occurrence with T11SS OMPs may hint at a role for the latter in molybdenum acquisition. The HrpC-like subcluster includes genes predicted to encode redox enzymes, such as formate dehydrogenase (48/81) and NADPH:quinone oxidoreductase (12/81), tRNA synthases and modification systems (30/81), and regulatory proteins including TetR family regulators, FaeA family regulators, FecR, and FecI (32/81). Homologs in the Hpl-like subcluster had few unifying co-occurrences, however many co-occurred with tRNA synthases/modification systems (14/36), specifically selenocysteine tRNA synthases (6/36) (Fig. 2B). Overall, network analysis of hemophilin homologs indicate that the family is diversifying into distinct subfamilies independent of their taxonomic lineage, and that those subfamilies are genomically associated with metal-related cellular activities such as ion uptake, metal responsive regulatory proteins, and metal dependent enzymes.

### Hemophilin cargo proteins have specificity for their cognate T11SS

Hemophilin homologs in each of the network subfamilies are predicted to be secreted by a T11SS, and many were encoded adjacent to a T11SS OMP. We considered the possibility that co-diversification of hemophilins and their T11SS OMPs has resulted in specificity between cognate pairs. To test the extent of specificity between hemophilin cargo proteins and their paired TXISS OMP secretor, and to further establish the role of TXISS OMPs in secretion of hemophilin cargo, we conducted secretion assays on different cognate or non-cognate pairs, chosen from each of the different subclusters of the hemophilin family network (except for the Plant/Environmental subcluster): HrpB_X.nem_/HrpC from *Xenorhabdus nematophila*, HrpB_H.haem_/Hpl from *Haemophilus haemolyticus*, HsmA/HphA from *A. baumannii*, and CrpB/CrpC from *Xenorhabdus cabanillasii* were cloned into pETDuet-1 based expression vectors to perform co-expression and secretion experiments in *E. coli* BL21 C43. Additionally, plasmids were constructed to co-express HrpB_X.nem_ alongside the non-cognate hemophilin homologs Hpl, HphA, and CrpC. Western blotting of supernatant and cellular lysates was performed to monitor cargo localization to the extracellular milieu. Extracellular HrpC and HphA both ran as single protein bands somewhat larger than would be predicted from mature protein sequence alone, with HrpC appearing at ∼30 kD (∼24 kD expected) and HrpA appearing at 29 kD (∼25 kD expected) (Supplemental file 2: sheets 2 and 4) [27]. In both cases this increase in apparent size is too large to be explained by the presence of the signal peptide, though this larger apparent size is consistent with our previous studies of HrpC [12]. Extracellular Hpl and CrpC were both observed running as two different bands. Hpl had a predominant band at ∼32 kD and a minor one at the expected size of ∼26 kD, while CrpC had a predominant band at ∼19 kD and a minor band at the predicted size of ∼27 kD (Supplemental file 2: sheets 3 and 5). Interestingly, the extracellular ∼19-kD CrpC band was absent or reduced when the protein was expressed alone or with the non-cognate HrpB_X.nem_, suggesting potential protein modification (e.g., cleavage) prior to, or during, secretion by its cognate T11SS. For enumeration of Hpl and CrpC localization we opted to sum both protein bands to fairly represent all of the secreted protein.

Co-expression of hemophilin homologs with a cognate T11SS protein always significantly increased the concentration of cargo protein found in the supernatant, though the level of secretion varied greatly among T11SS proteins. At one hour after induction, the presence relative to the absence of the cognate T11SS OMP increased the average level of extracellular hemophilin homolog protein (HrpB_X.nem_/HrpC: 34.2-fold; HrpB_H.haem_/Hpl: 59.3-fold, HsmA/HphA: 4.6-fold; CrpB/CrpC: 56.9-fold) (Fig. 3; Supplemental File 2). HrpB_X.nem_ significantly increased (4.8-fold) the average extracellular levels of the non-cognate cargo Hpl, though it was significantly less effective at doing so than the cognate T11SS HrpB_H.haem_. HrpB_X.nem_ did not significantly impact average extracellular levels of HphA, (1.1 fold), or CrpC (1.7 fold).

**Figure 3.**
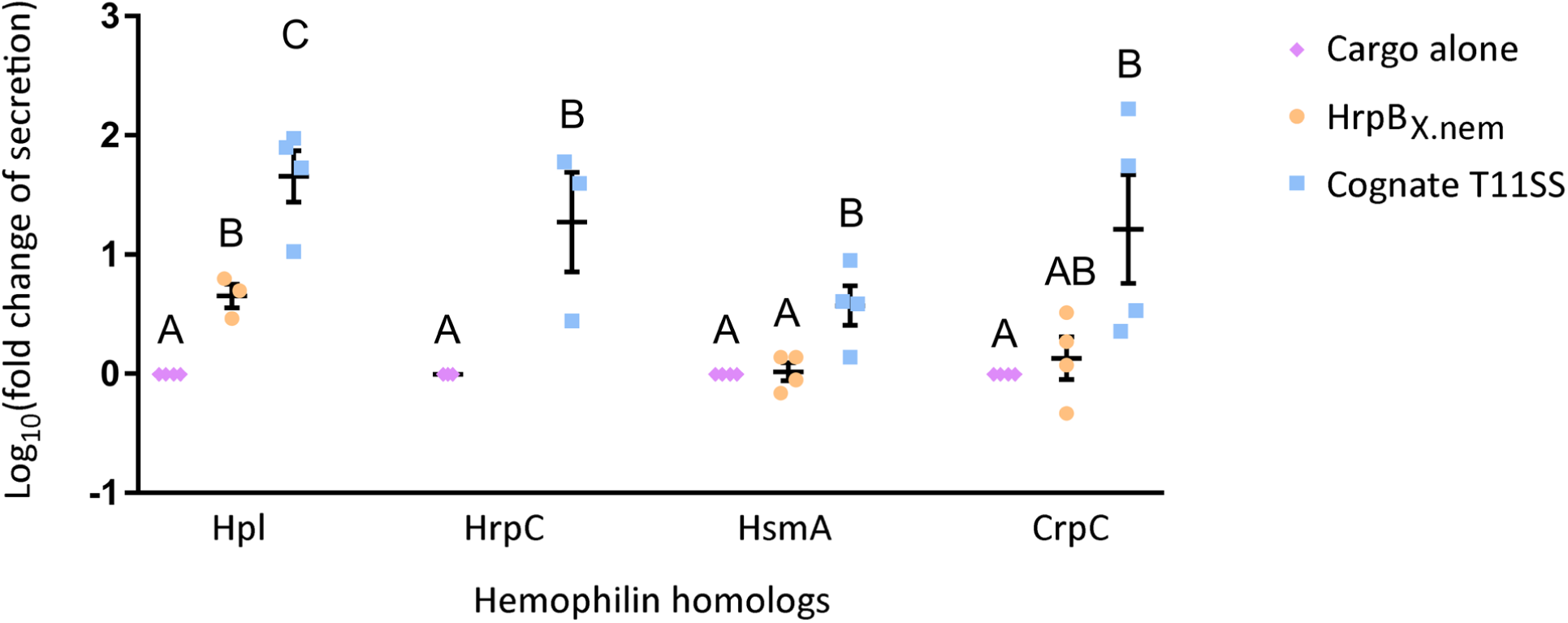
Secretion of hemophilin proteins by their cognate and non-cognate T11SS proteins. Each cargo protein was expressed in *Escherichia coli* in isolation (cargo alone; pink diamonds) or co-expressed in with their cognate T11SS (blue squares) or with non-cognate HrpB_X.nem_ (orange circles). Supernatant proteins were precipitated and quantified via immuno-blotting with anti-FLAG antibody to detect extracellular cargo proteins. Fold change of secretion was determined by dividing the amount of extracellular cargo detected in co-expression treatments by the amount seen in the respective cargo-alone treatment. Data were transformed with a log_10_ function prior to performing a Tukey’s HSD test for each hemophilin homolog. Letters indicate significance groups. All four T11SS proteins assayed significantly increased the amount of cognate cargo that was secreted into the extracellular milieu. The non-cognate T11SS HrpB_X.nem_ significantly increased secretion of Hpl but less effectively than the cognate T11SS of Hpl. HrpB_X.nem_ did not significantly affect the secretion of HsmA or CrpC.

### The hemophilin C-terminal β-barrel domain contributes to specificity for T11SS

The data described above indicates that individual hemophilin family cargo proteins and their cognate T11SS OMPs have specificity for each other. Published literature has implicated the C-terminal β-barrel domain of cargo proteins in directing secretion by T11SS OMPs [11]. To assess the role of the hemophilin C-terminal domain on T11SS specificity, two chimeric hemophilin cargo were engineered. The first chimeric cargo had the N-terminal handle domain from HrpC and the C-terminal β-barrel domain from Hpl (henceforth HrpC-Hpl), while the second had the N-terminal handle from Hpl and the C-terminal β-barrel domain from HrpC (henceforth Hpl-HrpC). pETDuet-1 constructs were assembled to independently co-express HrpB_H.haem_ and HrpB_X.nem_ alongside both chimeric cargo proteins (Fig. 4A) and Western blotting of supernatant and cellular lysates was again performed to monitor cargo secretion (Supplemental file 2: sheet 6). The HrpC-Hpl chimera ran as a single band at ∼27 kD, which is larger than the predicted size of ∼25 kD. The Hpl-HrpC chimera ran as two bands with apparent sizes of ∼33 kD and ∼24 kD, which straddle the expected size of ∼27 kD. Since Hpl-HrpC ran as two bands, similarly to Hpl, we opted to sum both bands for the purpose of enumeration. Co-expression with HrpB_H.haem_ increased on average, the extracellular levels of its cognate cargo Hpl by 65.0-fold, the HrpC-Hpl chimera by 51.0-fold, and the Hpl-HrpC chimera by 7.8-fold, indicating that the presence of the cognate C-terminal domain of Hpl was optimal for secretion by HrpB_H.haem_. Co-expression with HrpB_X.nem_ increased on average, the extracellular levels of the cognate HrpC by 15.6-fold, the HrpC-Hpl chimera by 5.1-fold, and the Hpl-HrpC chimera by 8.1- fold on average (Fig. 4B; Supplemental File 2). Overall, both the presence of the C-terminal domain of the cognate cargo, relative to the non-cognate cargo, resulted in higher levels of secreted chimeric hemophilin homolog protein. This finding reinforces the concept that T11SS cargo are two-domain proteins: An N-terminal effector domain and a C-terminal β-barrel domain that directs cargo for T11SS-mediated secretion [28]. Our findings additionally implicate the C-terminal β-barrel domain in mediating specificity of cargo for particular T11SS OMPs.

**Figure 4.**
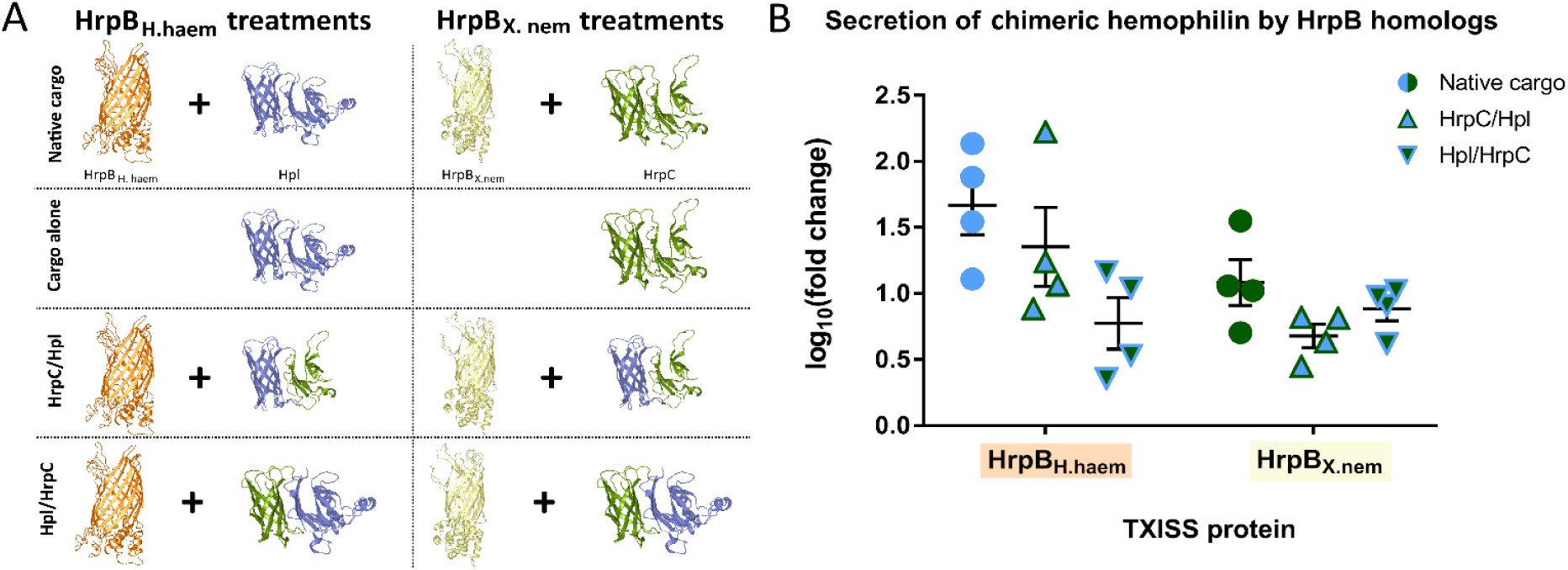
Secretion of domain-swapped chimeric hemophilin proteins by HrpB_X.nem_ and HrpB_H.haem_. **A)** A pictorial depiction of the experimental treatments used to assess secretion of chimeric hemophilin cargo proteins by HrpB T11SS secretors from *H. haemolyticus* (HrpB_H.haem_; orange) and *X. nematophila* (HrpB_X.nem_; yellow). Each T11SS protein was co-expressed alongside its cognate cargo protein (Hpl [blue] and HrpC [green], respectively), and two chimeric cargo proteins (HrpC/Hpl and Hlp/HrpC) generated by swapping the two domains (N-terminal effector/C-terminal barrel) of Hpl and HrpC. Fold change of secretion was determined by immunoblotting with anti-FLAG antibodies and dividing the amount of extracellular cargo in detected co-expression treatments by the amount seen in the respective cargo-only treatment. **B)** Secretion data were made normal with a log_10_ transformation. Both chimeric hemophilin proteins were preferentially secreted by the T11SS that was cognate to the C-terminal barrel domain they contained. However, neither chimeric protein (triangles) was as effectively secreted as the native (non-chimeric; circles) hemophilin proteins.

### Hemophilin homologs Hpl, HphA, and HrpC, but not CrpC bind heme

To begin to investigate if the heme ligand binding properties of Hpl and HphA are conserved in other hemophilin homologs, Hpl from *H. haemolyticus*, HrpC from *X. nematophila*, HphA from *A. baumannii*, and CrpC from *X. cabanillasii* were expressed without signal peptides in the cytoplasm of *E. coli* and purified. As expected, Hpl from *H. haemolyticus* was recovered from *E. coli* cytoplasm as an approximately 50:50 mix of heme-bound and heme-free protein, with the level of heme saturation likely reflecting competition for heme binding and limitations of heme biosynthesis *in vivo* [18]. Preparations of HrpC (*X. nematophila*) and HphA (*A. baumannii*) had a brownish appearance and an absorbance peak at ∼413 nm, possibly indicating the presence of sub-saturating levels of a porphyrin ligand, whereas CrpC (*X. cabanillasii)* was colorless with no peaks in the visible absorption spectrum, indicating the lack of a porphyrin. Heme-free (apo-protein) preparations of Hpl, HphA, and HrpC were produced by acid- acetone extract and reversed-phase HPLC, and heme-binding activities were investigated by titration (Fig. 5; Table 1). Large changes in the UV-visible spectrum of hemin occurred upon titration with *H. haemolyticus* Hpl, *A. baumannii* HphA or *X. nematophila* HrpC (Fig. 5A–C). Similarities in the Soret (412–414 nm) and Q-band regions (500–600 nm) between HrpC and HphA suggest that the heme coordination structure of HrpC is similar to that in HphA [13]. The binding curves for Hpl, HphA, and HrpC (Fig. 6A) yielded *K*_d_ values of 9, 7 and 20 nM respectively; however, the curves were close to linear, indicating that binding might be too strong to reliably extract binding constants. Using simulated data with added Gaussian noise we determined that data generated with *K*_d_ values < 15 nM produced fits that did not differ significantly (at P ≤ 0.05 by F test) from fits where the *K*_d_ parameter was fixed at an extreme low value. Thus, we suggest that Hpl and HphA bind heme with *K*_d_ values ≤ 15 nM and HrpC binds with *K*_d_ = 20 (10–50) µM. In contrast, spectral changes upon addition of CrpC to heme were more gradual (Fig. 5D) and were fit with a one-to-one binding model with apparent *K*_d_ ∼5 μM (Fig. 6A). These values are similar to the binding affinity we determined for BSA with apparent K_d_ ∼2 μM (Fig. S2).

**Figure 5.**
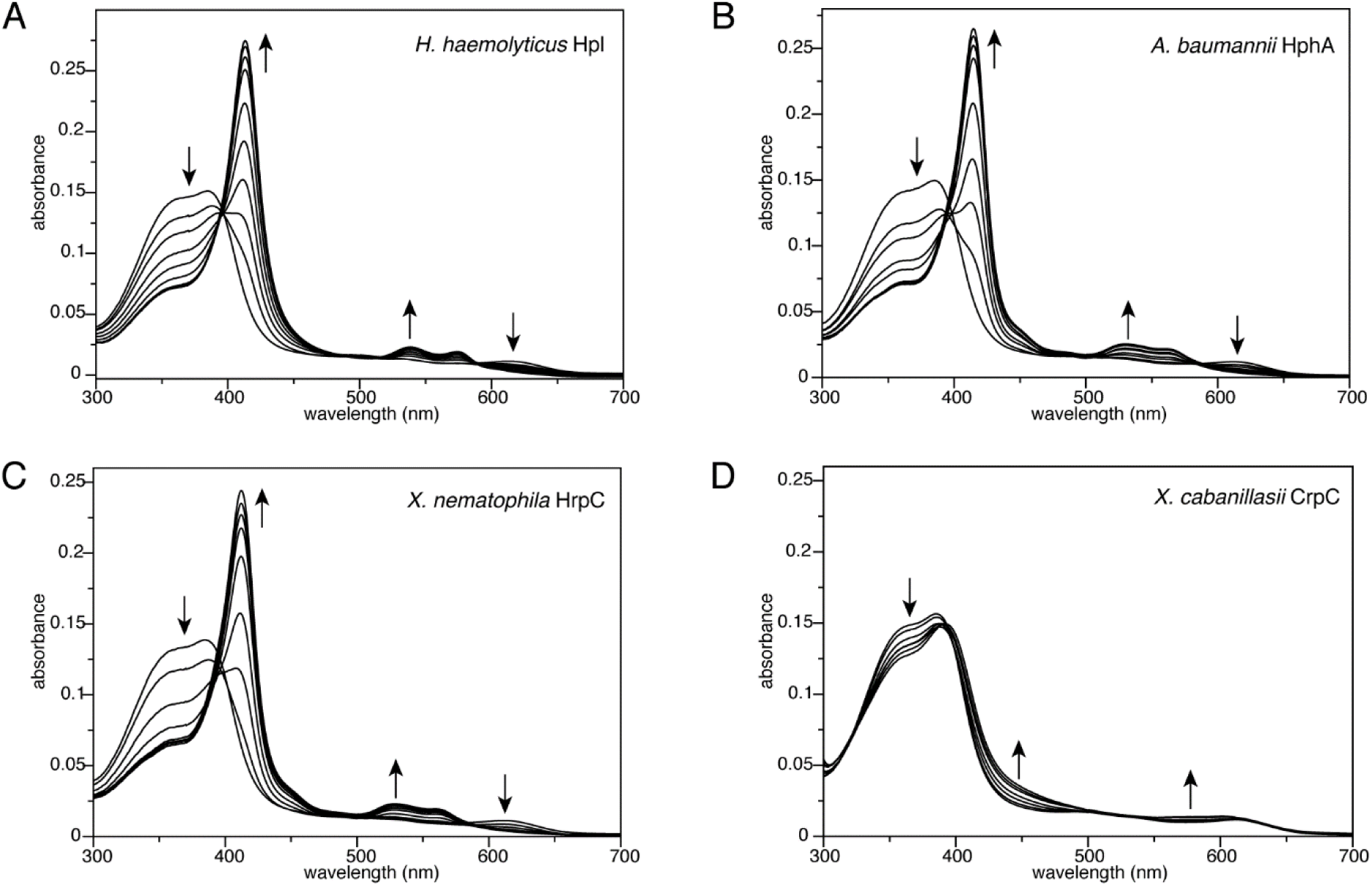
Titration of hemophilin homologues with hemin. Titrations of hemophilin proteins into hemin (Fe(III)-PPIX) solution (2.5 µM hemin in 20 mM Tris-HCl, pH 8.0 at 21°C) were monitored by UV-visible absorption spectroscopy. **A)** Spectra recorded after addition of *H. haemolyticus* Hpl to a final concentration in the range 0.3–4.5 µM. **B)** Spectra recorded after addition of *A. baumannii* HphA to a final concentration in the range 0.4–3.8 µM. **C)** Spectra with addition of *X. nematophila* HrpC in the concentration range 0.2–5.4 µM. **D)** Spectra with addition of *X. cabanillasii* CrpC in the concentration range 0.3–9.1 µM. Arrows indicate the direction of spectral changes.

**Figure 6.**
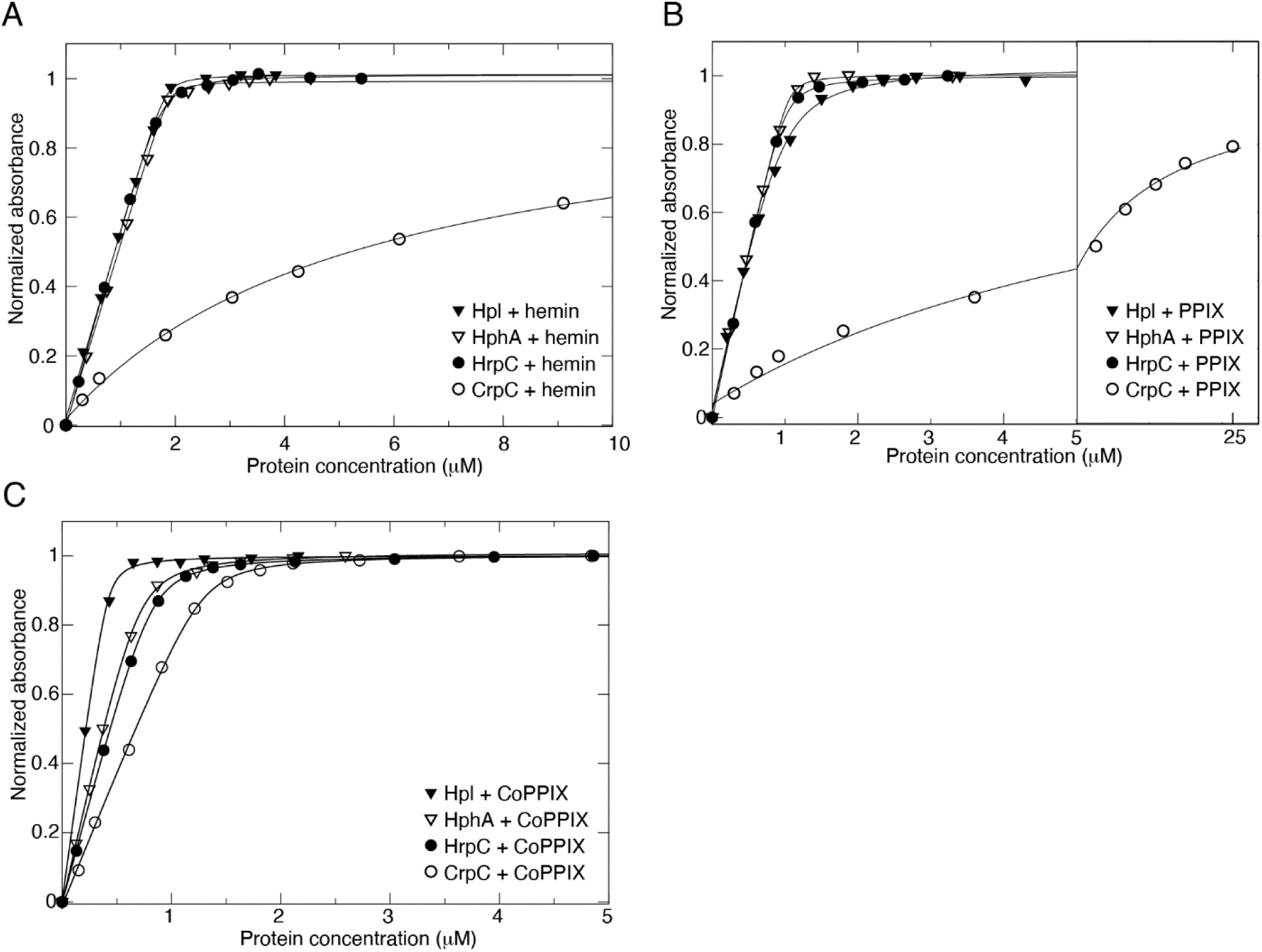
Binding isotherms for hemophilin homologs binding to porphyrins. Binding isotherms were generated from absorption data (symbols) by fitting to peak wavelength changes using a 1:1 binding model accounting for ligand depletion (lines). **A)** Binding isotherms for Hpl, HphA, HrpC or CrpC with hemin, generated from data in Figure 5. **B)** Binding isotherms for Hpl, HphA, HrpC or CrpC with PPIX, generated from data in Supplementary Figure 2. **C)** Binding isotherms for Hpl, HphA, HrpC or CrpC with Co(III)PPIX, generated from data in Supplementary Figure 4.

**Table 1.** Relative porphyrin binding affinity of hemophilin homologs

| Protein | Hemin<br>$K_d$ (nM)<sup>†</sup> | Zn(II)-PPIX<br>$IC_{50}$ (nM)<sup>*</sup> | Zn(II)-PPIX<br>$K_d$ (nM)<sup>†</sup> | Zn(II)-PPIX<br>$K_d$ (nM)<sup>‡</sup> | PPIX<br>$K_d$ (nM)<sup>†</sup> | Co(III)-PPIX<br>$K_d$ (nM)<sup>†</sup> |
| --- | --- | --- | --- | --- | --- | --- |
| Hh Hpl | ≤ 15 | 10 (1 – 20) | N/A | 0.2 | 70 (30 – 140) | ≤ 15 |
| Ab HphA | ≤ 15 | 20 (2 – 50) | N/A | 0.4 | ≤ 15 | 30 (10 – 50) |
| Xn HrpC | 20 (10 – 50) | 110 (70 – 180) | N/A | 2.2 | 20 (10 – 30) | 24 (20 – 29) |
| Xc CrpC | 5 (1–9) × 10<sup>3</sup> | N/A | 3 (2 – 4) × 10<sup>2</sup> | N/A | 7 (3–11) × 10<sup>3</sup> | 24 (21 – 26) |
| BSA | 2 (1–5) × 10<sup>3</sup> | N/A | 3 (2 – 4) × 10<sup>2</sup> | N/A | 11 (6–14) × 10<sup>3</sup> | 70 (50 – 80) |

Values in brackets are 95% confidence intervals
\* Calculated from fluorescence measurements in competition with BSA
† Calculated from absorbance data
‡ Calculated from $IC_{50}$ values according to Cheng and Prusoff 1973, using measured $K_d = 0.3$ μM for BSA binding Zn(II)PPIX and [BSA] = 15 μM

To further distinguish between the porphyrin binding affinities of Hpl, HphA, and HrpC, we assayed binding to Zn(II)PPIX, a fluorescent heme analog, in the presence of competition from excess BSA (Fig. 7). We reasoned that fluorescence detection and competition binding would shift the useful detection range to a high-affinity regime, compared to the previous absorption design. These experiments (Fig. 7) yielded IC_50_ values for hemophilin binding Zn(II)- PPIX in competition with BSA as shown in Table 1, together with calculated affinities based the affinity of BSA for Zn(II)PPIX. The IC_50_ values for Hpl and HphA did not differ within error of the measurements, suggesting that Hpl and HphA bind Zn(II)-PPIX with similar affinity. In comparison, the IC_50_ value for *X. nematophila* HrpC was approximately 5–10 fold higher, indicating weaker binding to Zn(II)-PPIX. Using the same fluorescence competition assay, the interaction of CrpC with Zn(II)-PPIX was undetectable, consistent with the finding that CrpC and BSA have similar propensities to bind heme. In summary, the above results suggest that heme binding affinities of the hemophilin homologs proceed from higher to lower affinity in the order Hpl ≍ HphA > HrpC >> CrpC.

**Figure 7.**
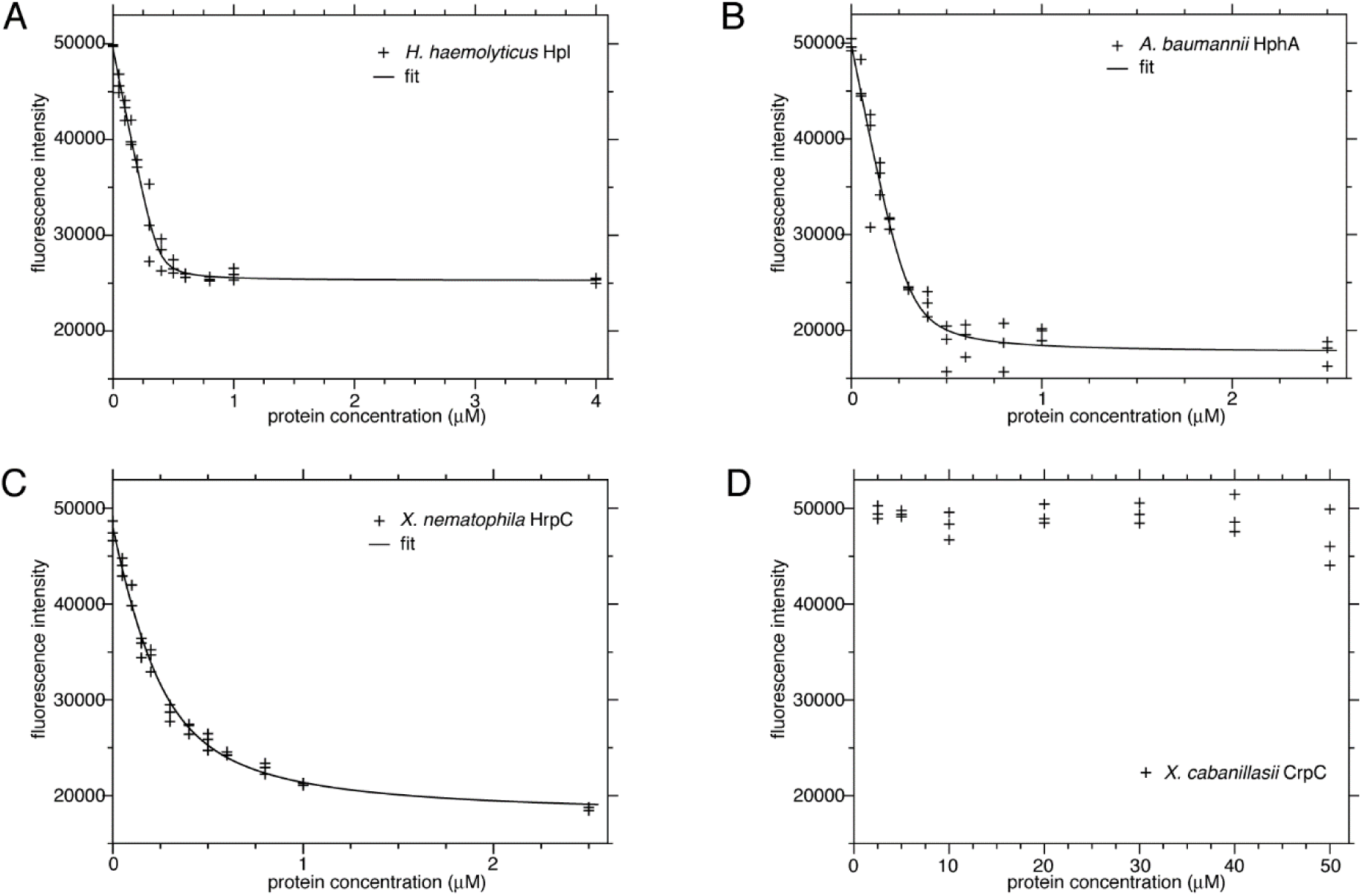
Titration of hemophilin homologues with Zn(II)PPIX. Binding of hemophilin homologues was monitored by changes in fluorescence intensity of the Zn(II)PPIX ligand (prepared in 20 mM Tris-HCl, pH 8.0 at 21°C) in the presence of excess BSA (15 µM). Data points (+) were fit (solid lines) with a 1:1 binding model accounting for ligand depletion. **A–C)** Titrations of Zn(II)PPIX (0.5 µM) with Hpl, HphA or HrpC. **D)** Titration of CrpC into Zn(II)PPIX (1.0 µM).

To investigate the importance of the porphyrin metal to hemophilin binding, we performed spectroscopic titrations with unmetalated PPIX. Because the CrpC gene is frequently located close to cobalt/molybdenum associated genes, we also screened for binding to Co(III)PPIX. Titration, monitored by UV-visible spectroscopy, indicated that Hpl, HphA and HrpC bound PPIX with nM affinities, whereas CrpC binding was much weaker (*K*_d_ ∼7 µM), in a pattern (affinity HphA > Hpl ≍ HrpC >> CrpC) similar to that seen for a hemin ligand (Fig. 6B and S3). A different pattern was seen with Co(III)-PPIX. Titration experiments with Co(III)-PPIX indicated affinities ranked in the order Hpl > HphA ≍ HrpC ≍ CrpC, corresponding to a dramatic increase in the relative affinity of CrpC for Co(III)PPIX compared to other porphyrins we tested (Fig. 6C and S4). To understand the potential significance of this we looked at the interaction of BSA with Co(III)-PPIX. We found that Co(III)-PPIX, and to a lesser degree Zn(II)-PPIX, bound more strongly to BSA than did PPIX or hemin, suggesting that the Co(III) and Zn(II) metals might contribute to binding interactions that are not available to Fe(III). We tested several other porphyrin-related molecules, including coproporphyrin III, biliverdin and cobalamin and found no evidence for CrpC binding (Fig. S5). These results suggest that Hpl, HphA, and HrpC might effectively scavenge metalated or unmetalated porphyrins from the environment. *X. nematophila* HrpC appeared to bind metalated and unmetalated porphyrin with similar affinities (10–20 nM). In contrast, CrpC binds very weakly to PPIX and hemin. It is interesting to consider whether this weak binding could allow CrpC to preferentially bind cobalt porphyrins in a biologically setting.

### Hemophilin family proteins adopt a multi-domain structure characteristic of T11SS cargo

Our observations that hemophilin homologs vary in their binding capacities for porphyrins prompted us next to investigate the structure function relationships among them. The Hpl and HphA crystal structures [13,18] were compared against the protein structure database (DALI server [29]) and manually inspected within the protein fold classification databases (CATH [30] and SCOP [31]). We observed the expected C-terminal β-barrel domain structure (Fig. 8A and 8C) that appears to be characteristic of T11SS cargo proteins (CATH superfamily 2.40.160.90, SCOP superfamily 3002098) [18]. More limited similarity was detected between the N-terminal ligand-binding domains of Hpl or HphA (Fig. 8B and 8D) and the N- terminal handle domain of TbpB proteins (CATH superfamilies 2.40.128.240/250; SCOP fold 2001281). This region also appears to be characteristic of T11SS cargo proteins, including Hpl and HphA, and forms a β-sheet (strands 1, 8, 7, 6, and 5 in Hpl and HphA) that packs against the β-barrel domain. The remainder of the N-terminal region of the Hpl and HphA are variable in structure compared to each other and the other known T11SS cargo proteins. Whilst the N- terminal domains of Hpl and HphA share features with the handle domain of TbpB, the insertion of one or more α-helical elements between β-strands 3 and 4, and a change in hydrogen-bond connectivity within β-strands 1–4 suggest that the hemophilin ligand binding domain belongs to a separate domain family, as a subgroup of the TbpB handle domain topology.

**Figure 8.**
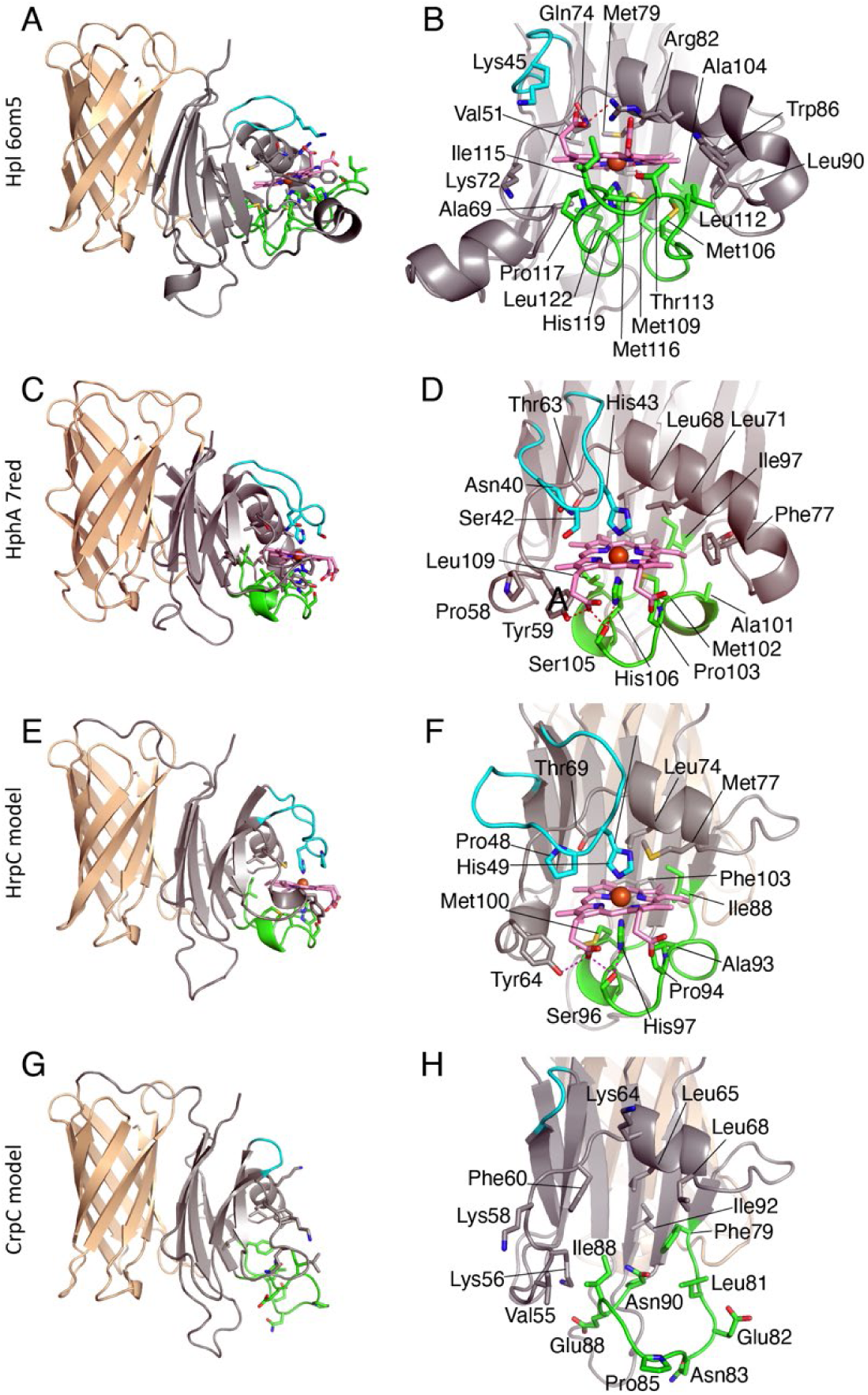
Structural modelling of HrpC and CrpC. **A)** Domain structure of Hpl (pdb 6om5) showing the N-terminal hemophilin ligand binding domain (grey, cyan, green) and C-terminal 8- stranded β-barrel domain (BBD; tan color). **B)** Heme binding site of Hpl showing structural elements that contribute to heme binding including the β2-β3 loop (cyan), β3-β4 α-helix that carries Arg82 and Trp86, and the β5-β6 loop (green) that carries the heme-coordinating His119. The heme porphyrin and central iron atom are shown in pink and orange, respectively. **C)** Domain structure HphA (pdb 7red) shown with the same coloring as A. **D)** Heme binding site of HphA colored as B. The heme iron is coordinated by two His side chains: His43 in the β2-β3 loop and His106 in the β4-β5 loop. **E-F)** Structural model of *X. nematophila* HrpC based on the alignment in Figure 10, modelled with bound heme. The heme binding site has features similar to HphA, including bis-histidyl heme coordination. **G-H)** Modelled structure of *X. cabanillasii* CrpC without a heme ligand. The precise conformation of the β5-β6 loop cannot be modelled by threading due to the lack of a suitable template.

We also considered the predicted structures of HrpC and CrpC as representatives of the two other subclusters of hemophilin family proteins. Overall, sequence similarity between hemophilins is higher in the β-barrel domain (40% identity between Hpl and HrpC; 36% identity between Hpl and CrpC) it is lower in the N-terminal domain (24% identity between Hpl and HrpC; 20% identity between Hpl and CrpC). We made structural models using the program MODELLER [32] with the crystal structures of Hpl and HphA [13,18] as templates (see methods) and found that HrpC (Fig. 8E-F) and CrpC (Fig. 8G-H) are predicted to adopt a similar overall structure as Hpl and HphA.

### Hemophilin homologs share a signature N-terminal ligand binding domain

We next compared features of the known or predicted N-terminal ligand binding domain folds of the hemophilin subcluster representatives, including from the Plant/Environmental subcluster. Alignments were used to model the structures of representative members of each subcluster, including *X. nematophila* HrpC (Fig. 8E-F) and *X. cabanillasii* CrpC (Fig. 8G-H) for which we had measured ligand binding. The ligand-binding domains of Hpl and HphA have a conserved hydrophobic core that was preserved in alignments with HrpC and CrpC, as well as other members of each subcluster, including the Plant/Environmental subcluster (e.g., *Lysobacter enzymogenes* ALN55974 and *Stenotrophomonas maltophila* KUJ02124) (Fig. 9). We also looked for conserved pair-wise contacts between side chains that are distant in the primary sequence and that might provide additional confidence in modelling local structures. For example, we identified a DXNG[V/I] motif corresponding to a β-hairpin in the ligand binding domain of *H. haemolyticus* Hpl that makes a bifurcated hydrogen bond with a Tyr side chain in the β-barrel domain (Fig. 10A) and is conserved in Hpl-like, HrpC-like and CrpC-like sequences, but is absent in the HphA-like cluster (Fig. 9). With respect to this motif, the Plant/Environmental subcluster comprises two subsets, one containing the DXNG motif (represented by ALN55974 from *L. enzymogenes*), and one lacking the DXNG motif and with a divergent sequence/structure in β4-β5 hairpin (represented by KUJ02124 from *S. maltophilia*) Fig. 9). Several groups of sequences in the Hpl-like subcluster could not be aligned or modelled against the hemophilin domain due to greater sequence (and therefore presumably structural) diversity) within this subcluster. With the exception of this group, our alignment and structural motif analyses indicate that hemophilin homologs (e.g., HrpC, CrpC, HphA, Plant/Environmental subclusters, and a subset of the Hpl subcluster) have an N-terminal domain with the same topology as the ligand-binding domains of *H. haemolyticus* Hpl and *A. baumannii* HphA and that this hemophilin ligand binding domain is the signature feature of an extensive family of hemophilin homologs.

**Figure 9.**
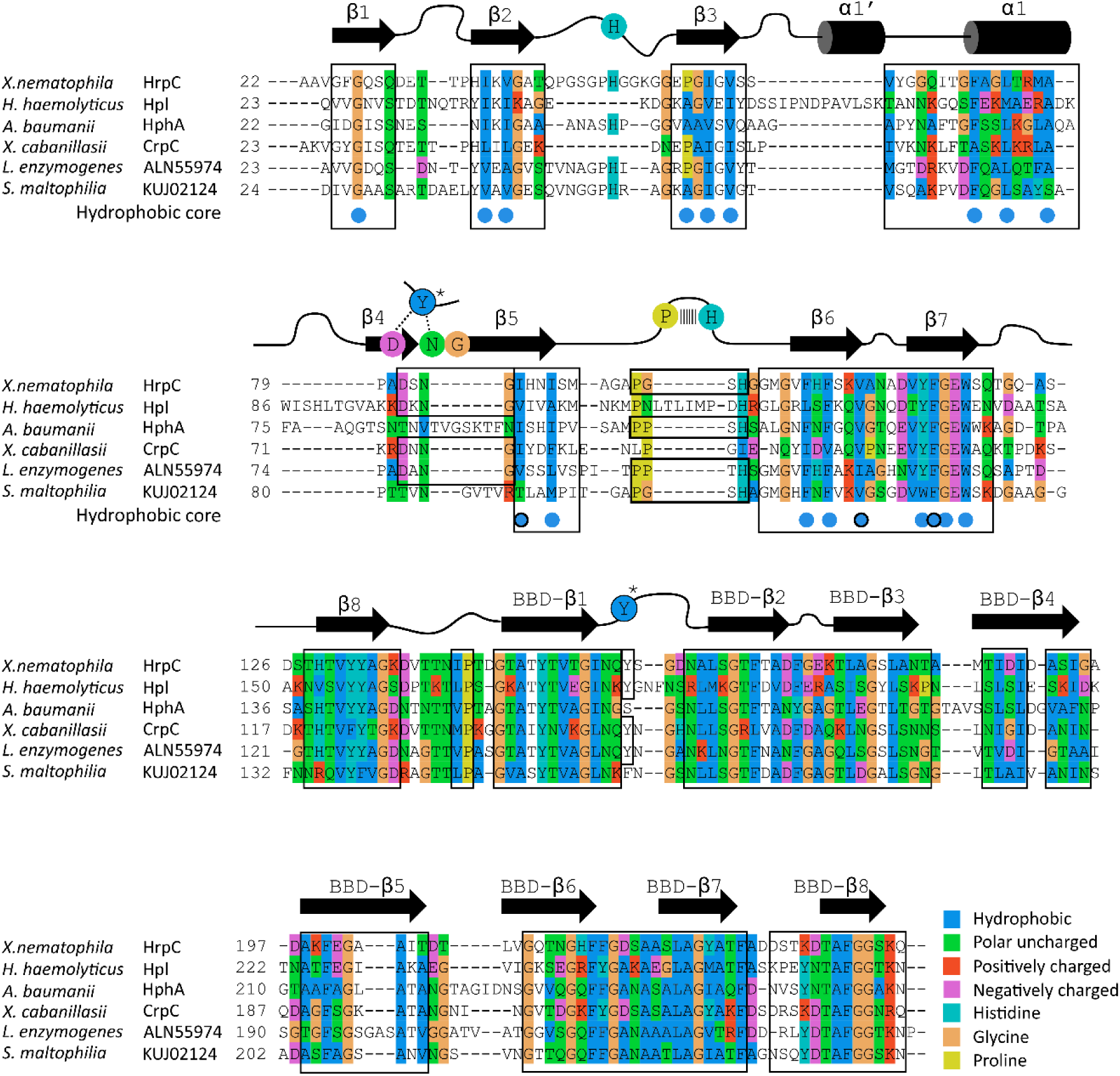
Aligned sequences of representative members from hemophilin subclusters defined by sequence similarity network analysis. The alignment is a composite of information including structure-based alignment of *H. haemolyticus* Hpl and *A. baumannii* HphA sequences and sequence-based alignments using the full membership of each hemophilin subcluster output from network analysis. Only representative members of the HrpC-like, CrpC-like and Plant/Environmental clusters used in modelling (see Figure 8 and supplementary Figure S6) are shown in the figure. The approximate position of secondary structure elements is displayed in cartoon, with residues of the DXNG and PX[S/T]H motifs (see Figure 10) shown in colored circles. Boxes indicate regions where query sequences could be modelled on one or both structure templates. Residues of the conserved hydrophobic core of the N-terminal hemophilin domain, or interdomain contact, are marked with blue circles without or with a black outline, respectively.

**Figure 10.**
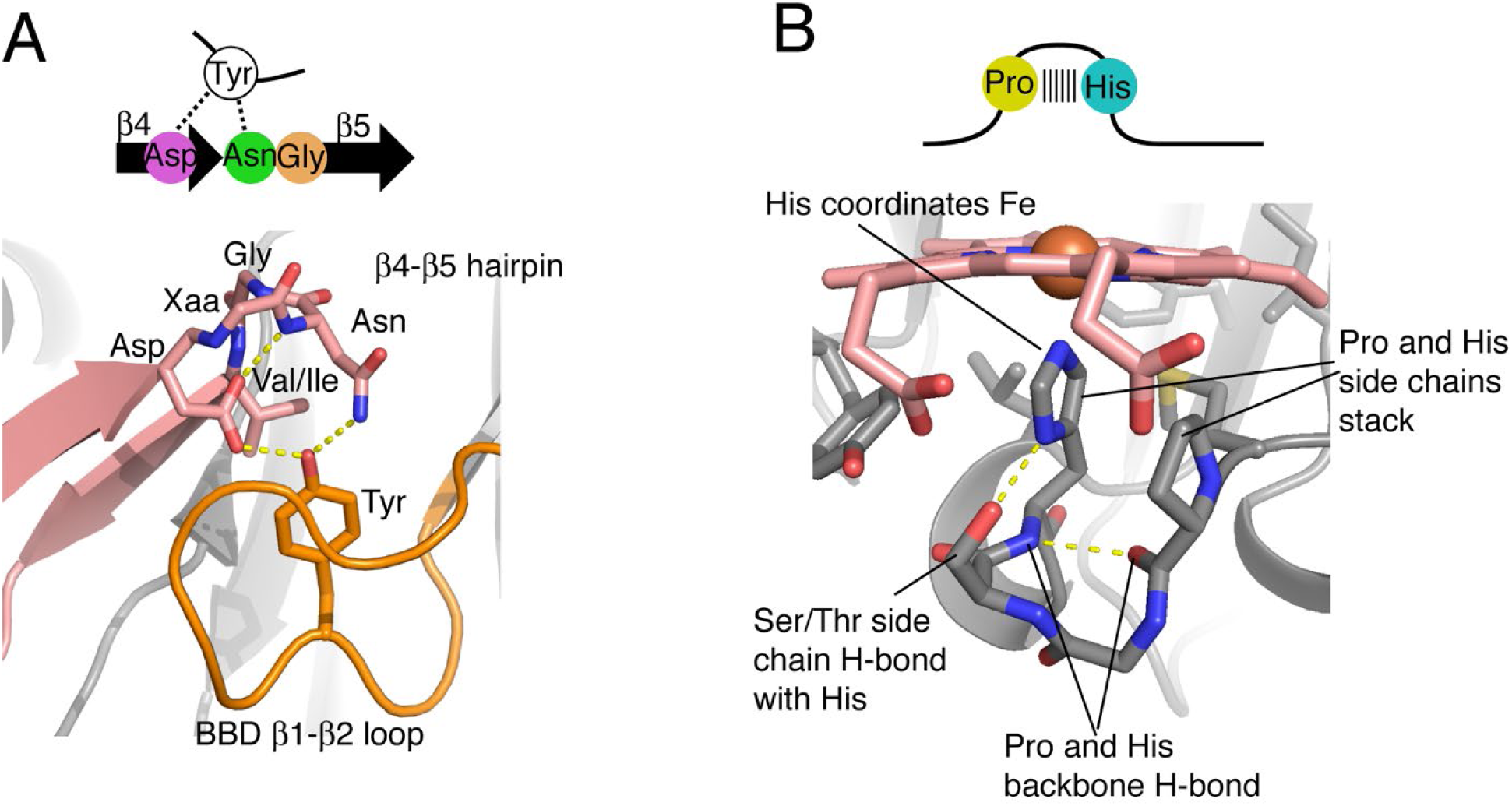
Conserved sequence-structure motifs. **A)** DXNG[V/I] motif structure in the β4-β5 hairpin. **B)** PX[S/T]H motif incorporating the heme-ligating His in the β5-β6 loop.

### Two distinct heme binding domains occur among hemophilin homologs

Having identified another hemophilin homolog capable of coordinating heme, we used models to investigate possible structural reasons for common and distinct ligand binding properties of hemophilin homologs. The available crystal structures of *H. haemolyticus* Hpl and *A. baumannii* HphA, both of which bind heme, show that residues in loops β2-β3, β3-β4 (including the β3-β4 α-helix) and β5-β6 contribute majority of heme-binding contacts and that the mode of heme binding in Hpl and HphA differs in the relative contributions of these loops and in the heme-iron coordination structure (Fig. 8B and 8D). Of particular interest, heme in Hpl is coordinated through a single His side chain in the β5-β6 loop, whereas heme in HphA is coordinated by two His side chains, one from the β5-β6 loop and a second donated by the β2-β3 loop.

The bis-histidyl heme coordination observed for HphA appears to be ubiquitous in the HphA subcluster, based on the conservation of the amino acid sequence of this motif (data not shown). A subset of sequences within the Hpl subcluster had >65% identity to *H. haemolyticus* Hpl, and these are predicted to all share the same single histidinyl coordination heme binding mode. However, the heme coordination pattern for the remaining members of the Hpl subcluster is unclear due to the high level of sequence/structure diversity in this group. Molecular models show that HrpC subcluster and Plant/Environmental subcluster proteins contain His side chains in both the β2-β3 loop and β5-β6 loops, and modelling in the presence of a heme ligand indicates that the position of these His residues is compatible with bis-histidyl heme coordination similar to HphA (Fig. 8F and Fig. S6). Other structural similarities among HphA, HrpC, and Plant/Environmental subclusters, but not for the Hpl subcluster, exist in the β2-β3 loop, including a Gly, Ser, Pro rich sequence, and similar loop length (Fig. 9). The mechanism of heme binding and release by *A. baumannii* HphA is proposed to involve unfolding of the β2-β3 loop [13], and if so, this mechanism is likely to be conserved in hemophilin homologs with this motif. The second heme-coordinating His of HphA occurs in the β5-β6 loop, which contains a conserved PX[S/T]H sequence motif.

Like the β2-β3 loop, the PX[S/T]H motif of the β5-β6 loop is conserved across the HphA, HrpC and Plant/Environmental clusters (Fig. 9, Fig. 10B). In *A. baumannii* HphA, the His side chain of this motif is packed against the pyrrolidine ring of Pro and makes a H-bond with the Ser side chain (substitution of Thr for Ser is expected to preserve this interaction). The carbonyl oxygen of Pro also accepts a H-bond from the backbone amide of His. These interactions are expected to stabilize the orientation of the heme-ligating His, and conservation of this motif in HrpC and Plant/Environmental cluster hemophilin homologs further implies that the mode of heme binding in HphA, HrpC and Plant/Environmental proteins is similar. The PX[S/T]H motif is absent in Hpl, which has a substantially longer β5-β6 loop (20 residues in Hpl vs 14 residues in HphA) and more completely covers the face of the porphyrin.

### The N-terminal ligand binding domain regions of hemophilin homologs are diversifying

In addition to iron-ligating side chains, other amino acids contribute to ligand binding through non-covalent interactions with apolar and polar sites on the porphyrin. Models of *X. nematophila* HrpC show multiple hydrophobic contacts with the porphyrin ring contributed by side chains in the β3-β4 α-helix, β5-β6 loop and underlying β sheet, and a hydrogen bond network around the charged porphyrin 17-propionate comprising Tyr64, Ser96 and His97 side chains – features similar to the HphA heme site (Fig. 8, compare D, F). Differences include a shorter β3-β4 α-helix in HrpC than either HphA or Hpl, and a β4-β5 hairpin in HrpC that is more similar to Hpl than to HphA. The pyrrole rings B and C appear to be more exposed to solvent in HrpC than in either Hpl or HphA, which may partly explain a relatively lower affinity for the metalated porphyrins, hemin and Zn(II)PPIX, because hydration is expected to increases the rate of scission of the Fe-metal bond (although the lack of effect on Co(III)PPIX binding is not explained).

Our porphyrin binding assays revealed that *X. cabanillasii* CrpC does not bind heme. Consistent with this observation, while *X. cabanillasii* CrpC is predicted to adopt the hemophilin ligand binding domain fold, it lacks His or other residues that typically coordinate heme-iron, in either the β2-β3 or β5-β6 loops (Fig. 8G-H and Fig. S6). The β2-3 loop is reduced to a three-residue hairpin and is too small to contribute to a porphyrin pocket of the kind seen in HphA and HrpC. Neither does CrpC look like the Hpl heme-binding site, which employs a much longer β2-β3 α-helix and longer β5-β6 loop. Nevertheless, the distribution of apolar and polar side chains in Hpl/HphA that contact the porphyrin skeleton and ionizable propionates, respectively, is largely conserved in CrpC, although we cannot model the precise conformation of the β5-6 loop in CrpC due to lack of a suitable template. Thus, CrpC could potentially accommodate a porphyrin-like ligand in a more solvent exposed, and therefore presumably lower affinity, binding site. This *X. cabanillasii* distinct non-histidine ligand binding domain is shared among the other CrpC-like subcluster homologs, except for a small number of sequences that have a large deletion in the hemophilin domain that removes the porphyrin binding site altogether.

## Discussion

A cornerstone of microbial existence is the extracellular deployment of metal-binding molecules, such as hemophilins which facilitate the competitive uptake of metals essential for cellular physiology. Here we investigated the structural features, secretion, and ligand binding of representative hemophilins, a family of proteins in five distinct sequence similarity subclusters encoded by Proteobacteria from human, animal, plant, built and free-living environments. We found that the entire hemophilin family shares common motifs present in the solved structures of the hemophilins *H. haemolyticus* Hpl and *A. baumannii* HphA [13,18]: a central β-sheet domain and a C-terminal β-barrel domain, both of which are previously described characteristics of T11SS cargo proteins [11,12]. In contrast, the N-terminal ligand binding region was conserved in some, but not all hemophilin homologs. We established that, like the hemophilins *X. nematophila* HrpC and *A. baumannii* HphA [12,33,34], two other hemophilin homologs, *H. haemolyticus* Hpl and *X. cabanillasii* CrpC rely on a T11SS secretor to reach the extracellular milieu. This establishes T11SS-dependence across all four hemophilin sequence subclusters tested, and indicates T11SS-dependent secretion is a hallmark of the entire family. Further, the results presented here support the concept that the β-sheet-β-barrel domains are an ancestral scaffold for T11SS-dependent secretion, onto which new effector variants are emerging through evolution of the N-terminal domain.

Despite the overall conservation of the core β-sheet-β-barrel motif among T11SS- dependent cargo, there must be inherent differences among these domains, since different cargo proteins show specificity for different T11SS secretors. To explore the underlying contributors to such specificity, we capitalized on the fact that each hemophilin homolog is associated with a cognate T11SS secretor, enabling different combinations of related cargo- secretor pairs to be assessed for secretion activity. We found that, despite sequence, structural, and functional similarity among hemophilin homologs and among their cognate T11SS secretors, individual cargo-secretor pairs have specificity for each other, indicating co-adaptation and co-diversification. These results establish that T11SS specificity is not shared between cargo, even between closely-related homologs, and that the efficacy or rate of secretion varies between T11SS/cargo pairings. By testing the secretion of hemophilin chimeras with non-native C-terminal β-barrel domains we determined that this domain is largely responsible for specificity of targeting to its cognate T11SS secretor. The effectiveness of the tested secretors (*H. haemolyticus* HrpB and *X. nematophila* HrpB) was highest for the cognate hemophilin cargo protein, followed by the chimeric hemophilin with the cognate C-terminal β- barrel domain. The chimeric hemophilin with the non-cognate C-terminal β-barrel domain was not secreted. Chimera secretion did not reach the same levels as native hemophilin cargo protein-T11SS secretor pairings, which may indicate that the conserved β-sheet, which was not substituted within the chimeric proteins, might also contribute to specificity. Alternatively, it may indicate that the chimeric proteins generated have compromised stability relative to the native cargo sequences that slows transposition or speeds up protein degradation.

The ability of the tested T11SS secretors to distinguish their cognate C-terminal domain is striking given that their cargo proteins, Hpl and HrpC are closely related in sequence (39% identity) and function, and reveals a previously underappreciated resolution of T11SS cargo selection. Further, we show here that when expressed in a common *E. coli* background under identical conditions, the T11SS secretors vary in their magnitude or efficacy. For example, HrpB increased extracellular Hpl by 59.3-fold while HsmA increased extracellular HphA 4.6-fold. These aspects of secretion specificity and efficacy are important to consider for potential T11SS bioengineering applications. For instance, our findings raise the possibility that novel T11SS- dependent cargo could be engineered for specific secretion by fusing novel N-terminal functional domains to the C-terminal domain from a characterized cargo protein. Similarly, Slam proteins, the subset of T11SS responsible for surface exposure of lipoproteins, have been proposed as a novel mechanism for surface presentation of immunogenic antigens as a potential vaccination strategy [35], and our discovery that different T11SS may have different levels of secretion efficacy could be exploited to fine-tune the levels of surface exposure or secretion. An important next step towards achieving this goal will be to identify the individual sequence and structural motifs responsible for the specificity between cargo and T11SS and any T11SS-dependent cargo modifications that may occur, to enable automatic annotation of T11SS cognate pairs and informed engineering of novel pairings.

The results presented here provide a comparative framework in which to consider the evolution and relatedness of T11SS cargo protein structures, which can be described as comprising three domains: an N-terminal variable effector domain, a mid-protein β-sheet domain, and a C-terminal β-barrel domain. The C-terminal β-barrel domain, comprising 8 β- strands arranged in a meander topology with a sheer value of 8, and a central core of hydrophobic side chains [18], is a unique and defining feature of T11SS cargo proteins that, as we have established here, contributes substantially to the recognition of cargo by the cognate T11SS secretor. The β-sheet domain is also universally present among known T11SS cargo. It is formed in large part by amino acids in the central portion of the protein sequence, and stacks against the β-barrel forming a physical barrier and bridge between it and the variable N-terminal effector domain. In some cargo, including hemophilins, there are integrated structural interactions between the variable N-terminal region and the β-sheet, with residues in the former contributing one or more β-strands to the β-sheet, and residues in the latter contributing key active site residues.

In the case of hemophilins, there is a conserved β-strand topology across the entire protein, comprising the effector strands (β2-β4), β-sheet strands (β1 and β5-8) and β-barrel strands (β9-18). Within this topology, a conserved hydrophobic core is formed by residues in the β1-β7 region, as are interfacial contacts between the β-sheet and β-barrel domains. Indeed, the most highly conserved sequence motif in hemophilins is a YFGEW pentapeptide that maps to β7 and contributes side chains to both the hemophilin hydrophobic core and to interdomain contacts. The relatedness of the hemophilins with other metal-related T11SS cargo is supported by the fact that this conserved hemophilin ligand binding domain (Hlb) shares a parent fold (SCOP fold 2001281) with the TbpB handle domain (SCOP family 4006246; TbpB handle domain-like) and with HpuA (SCOP family 4005058, hemoglobin receptor HphA). However, it is distinguished from these by differences in connectivity in the β1-β3 strands and inclusion of one or more α-helical elements in the β3-β4 loop. These results imply that β-strand topology and the hydrophobic core of the Hlb comprise an ancestral scaffold which has been elaborated on in evolution to bind a diverse range of ligands: heme in the case of Hpl, HphA, and HrpC, a variety of protein targets in the cases of HpuA, TbpB, and fHbp, and unknown ligands in the case of CrpC and NilC.

When considering active site residues and topology, the hemophilin homologs fell into three general classes: the Hpl type with a single histidinyl coordination binding site, the HphA/HrpC type, which included representatives from the Plant/Environmental subcluster, with a bi-histidinyl coordination binding site, and the CrpC type which lacked any histidines in the predicted active site. The *A. baumannii* HphA might be considered closer to a canonical heme-binding structure, whereas the Hpl-like subcluster appears to be more diverse in structure and arrangement of heme ligands.

In the HphA/HrpC type class, the heme is coordinated between two histidines, one in the β2-β3 loop and a second in the β5-β6 loop. In all members of this binding class, the β2-β3 loop is rich in Gly, Pro, and polar residues. This low complexity sequence suggests that the conformational heterogeneity that is seen in HphA may be a general feature; in heme-bound HphA the loop and heme-coordinating His cover one face of the porphyrin, whereas the same loop points out away from the binding site in the apo protein [13]. The structure of the β5-β6 loop does not differ between holo and apo structures of HphA and is predicted to be conserved across the HphA, HrpC and Plant/Environmental subclusters; a characteristic PX[S/T]H motif that features the second heme-ligating His side is stabilized by π-stacking and H-bonding interactions. A search of the protein database using RASMOT-3D [36] identified only one other heme binding protein (nine-heme cytochrome c, pdb 19hc) with a similar 3D arrangement of Pro, Ser, and heme-coordinating His side chains, but different primary sequence arrangement, thus the PX[S/T]H appears to be specific feature of the hemophilin domain. The modelled CrpC structure is highly similar to HrpC except that the β2-β3 loop is reduced to a hairpin and PX[S/T]H motif is gone, suggesting an evolutionary trajectory for loss of heme binding in CrpC.

Whereas *H. haemolyticus* Hpl clearly adopts the same topology as HphA, HrpC, Plant/Environmental, and CrpC subcluster representatives (with addition of a second helical segment in the β3-β4 loop of Hpl), the Hpl subcluster exhibits more extensive sequence (and likely structural) variability. One common feature within the Hpl subcluster is a D[R/K/S]NGV motif in the β4-β5 hairpin that is predicted to make conserved interactions at the interface between the β-sheet and the β-barrel domains. A similar motif appears in HrpC, CrpC, and a subset of the Plant/Environmental subcluster, but it is absent in HphA and the remaining Plant/Environmental cluster sequences, indicating that some structural features do not conform to a simple division between HphA-like and Hpl-like sequences. The heme-associated α-helix in both HphA and Hpl is longer than predicted in HrpC and Plant/Environmental cluster proteins, possibly contributing to higher heme affinity in HphA/Hpl by protecting a larger surface area of the porphyrin from solvent. Although most sequences within the Hpl subcluster carry His in the putative β5-6 loop, one group (sequence OFR67839 is one example) lacks this residue and other subgroups potentially have two His ligands.

The distinguishing active site characteristics among the hemophilin homologs contextualized our findings that purified hemophilin homologs displayed variable affinity for hemin (Fig. 5; Table 1). Intriguingly, this variable affinity for hemin may reflect differences in the ecological roles of the hemophilin homologs. Hpl and HphA, which have established roles in competing for heme under stringent conditions, had the highest heme affinity. *H. haemolyticus* Hpl is a heme chelating protein for nutritional heme uptake and can prevent the related organism, *H. influenzae*, from accessing Plant/Environmental heme [18]. This role for *H. haemolyticus* Hpl as a tool for competing against other organisms by sequestering a limiting nutrient (a form of nutritional immunity) would naturally favor the evolution of high hemin affinity. In a similar vein, *A. baumannii* HphA competes with mammalian host nutritional immunity for heme. In contrast to these two hemophilins, HrpC, encoded by *X. nematophila*, displayed lower heme binding affinity. This may be a reflection of the distinctive animal host niches occupied by *X. nematophila* relative to *H. haemolyticus* and *A. baumannii*. As a pathogen, *Xenorhabdus* can infect diverse insects, which are notoriously heme poor [37] and as a mutualist, it colonizes the intestinal tissues of its nematode host, which is a heme auxotroph [38,39]. The insect *Drosophila melanogaster* transferrin-1 binds heme with lower affinity than mammalian transferrin and is susceptible to low pH conditions [40]. Therefore, to access heme in an insect environment, *X. nematophila* HrpC may not need high affinity to overcome host chelation. In turn, lower affinity may offer a selective advantage by enabling resource sharing with its mutualistic nematode host, *S. carpocapsae,* with the resulting improved fitness benefitting both mutualistic partners. Dissociation of heme from HrpC would be essential for heme to be transferred to nematode iron chelating proteins. Entomopathogenic nematodes can live for months at a time without feeding while in their free-living infective stage. During this time they have a sealed intestine and no access to exogenous nutrients and they may rely on their *Xenorhabdus* symbionts to provide heme chaperoned by hemophilin [41]. While this study did not test the heme-binding affinity of hemophilin homologs from the Plant/Environmental subcluster, these homologs encode His residues in the β2-β3 loop and the β5-β6 loop like HphA and HrpC (Fig. S6), indicating the potential for heme-binding activity. Future studies on the binding affinities of these homologs may shed light on additional functions of hemophilins and their roles in facilitating bacterial lifestyles within plants, built environments, and soil ecosystems.

Our work has revealed that not all hemophilins have heme-binding activity, which opens the possibility that this family is diversifying and potentially gaining new ligand binding activities. CrpC, a member of the Cobalt/molybdenum associated subcluster of hemophilin homologs, did not bind heme, consistent with the fact that the predicted active site had neither the heme-coordinating His residues noted for the other hemophilin classes, nor any other typical heme ligands such as Cys, Met, or Tyr. There are residue types in the CrpC β5-β6 loop (Asn, Gln, Glu) and β3-β4 α-helix (Lys), that conceivably could act as metal ligands; a small number of protein crystal structures show Asn and Gln as a second heme ligand together with His [42] or Gln [43], respectively, and a Glu side chain [44] is in within coordinate bonding distance of heme iron in the terminal cytochrome bd oxidase, though it does not contribute substantially to heme binding affinity [45]. Lys residues occur in several algal globins, but only as a second Fe-ligand together with His [46]. However, none of these possibilities fit with the predicted structure of CrpC, which does not have His or Cys residues close to the β5-β6 loop. Thus the absence of a strong heme-iron ligand provides a structural explanation for the failure of CrpC to bind to heme or Zn(II)PPIX.

In this context, the binding of Co(III)PPIX to CrpC with comparable affinity to other hemophilins was unexpected. However, it is intriguing given the genomic context of the genes encoding CrpC and its T11SS partner, CrpB, which are found adjacent to an anaerobic B12/cobalamin biosynthesis locus in at least three strains of *Xenorhabdus* (Fig. 2B; Supplemental file 2), which hints at a role for the CrpC/CrpB cargo/secretion pair in cobalt or cobalamin binding. We found that the affinity of BSA for Co(III)PPIX was similarly about 2 orders of magnitude stronger than binding to unmetalated PPIX suggesting that Co(III)PPIX makes useful contacts with a wider range of side chains, perhaps those present in the CrpC active site, than does either heme or Zn(II)PPIX. CrpC binds to Co(III)PPIX with similar affinity to other hemophilin homologues, but has a dramatically lower affinity for heme and PPIX compared to other hemophilin homologues. Based on these data, we suggest that CrpC has evolved to lose a heme binding function while maintaining the ability to bind an alternative cobalt-metalated porphyrin-related ligand, such as an intermediate in cobalamin biosynthesis or metabolism. Overall, our findings suggest that bacteria can encode multiple members of the hemophilin family, both bona-fide hemophilins, such as HrpC, and paralogs like CrpC, that might have evolved new ligand binding activities by divesting themselves of their affinity for heme. An exciting avenue for the discovery of new metal-, or other ligand-binding proteins will be to examine the activities of additional members of the hemophilin family of proteins.

## Materials and Methods

### Sequence similarity network analysis

Protein sequence similarity networks were generated using the Enzyme Function Initiative-Enzyme Similarity Tool (EFI-EST) [22]. As an input we used the previously reported database of soluble TbpBBD cargo encoding genes which co-occurred with T11SS encoding genes [12]. EFI-EST performs an all-by-all BLAST of query sequences to assess relatedness, and then generates a network where each node represents a protein sequence, and the color of the edges indicates relatedness between nodes. A minimum alignment score of 35 was chosen to reduce total network edges enough to visualize protein subclusters. To simplify visualization proteins sharing ≥ 80% identity were compressed into representative nodes. Networks were visualized and interpreted using Cytoscape v3.7.1 [47] and Gephi v0.9.5 [48]. For the complete network of soluble TbpBBD domain proteins, nodes were organized with the Fruchterman-Reingold force-directed algorithm [49]. For the network containing only hemophilin family proteins, nodes were organized with the ForceAtlas2 algorithm for continuous force-directed arrangement [50].

### Bacterial culture conditions

All strains, plasmids, and primers utilized in this study are described in supplemental file 3. All cultures were grown in either glucose minimal media [33], LB stored in the dark to prevent formation of oxidative radicals (henceforth dark LB), or glucose minimal media supplemented with 1% dark LB. Plate based cultures were grown on either LB supplemented with pyruvate to prevent the formation of reactive oxygen radicals (henceforth LBP or glucose minimal plates [33]. For plasmid-based expression, chemically competent *Escherichia coli* strain BL21-DE3 (C43) were chosen for ease of transformation and their ability to non-toxically express membrane proteins [51,52]. Strains of *Escherichia coli* were grown at 37°C. Where appropriate, media was supplemented with ampicillin at a concentration of 150 μg/ml, chloramphenicol at 15 μg/ml or kanamycin at 50 μg/ml unless another concentration is stated. Protein expression was induced at the midlog point of bacterial growth via addition of isopropyl β-d-1-thiogalactopyranoside (henceforth IPTG) at a concentration of 0.5mM.

### Construction of T11SS and T11SS-dependent cargo expression plasmids

Expression plasmids for HrpC alone (HGB2531) and HrpBC co-expression (HGB2530) were previously generated and reported on [12]. FLAG-*hrpB* was amplified from pETDuet- 1/*hrpBC_X.nem_*using primers 1-2. *hpl*-FLAG and its adjacently encoded T11SS neighbor, FLAG- *hrpB_H.haem,_*were amplified from the purified genome of *Haemophilus haemolyticus* BW1 using primers 3-6. *crpC*-FLAG and its adjacently encoded T11SS neighbor, FLAG-*crpB*, were amplified from the purified genome of *Xenorhabdus cabanillasii* (HGB2490) using primers 7-10. *hphA* (ACJ40780.1) and *hsmA* (ACJ40781.1) were generated via gene synthesis by Genscript. To make cargo-only expression plasmids, pETDuet-1, *hpl*-FLAG, *crpC*-FLAG, and *hphA*-FLAG were digested with NcoI and NotI. Each cargo protein was independently ligated into MCS1 of pETDuet-1 via T4 DNA ligase, resulting in pETDuet-1/*hpl* (HGB2526), pETDuet-1/*crpC* (HGB2525), and pETDuet-1/*hphA* (HGB2532). Integration of each T11SS-dependent cargo was confirmed via digestion with NcoI and NotI as well as Sanger sequencing using primers 11-12 at the University of Tennessee (UT) Genomics Core.

To make T11SS/cargo co-expression plasmids, each of the above cargo-only expression plasmids were digested with KpnI and NdeI, alongside the PCR products for FLAG-*hrpB_H.haem,_* FLAG-*crpB*, and FLAG-*hsmA*. Each T11SS protein was then independently ligated into MCS2 of the plasmid containing its cognate cargo via T4 DNA ligase, resulting in pETDuet- 1/*hpl*/*hrpB_H.haem_* (HGB2523), pETDuet-1/*crpC*/*crpB* (HGB2524), and pETDuet-1/*hphA*/*hsmA* (HGB2533). Additionally, the PCR product for FLAG-*hrpB* from *X. nematophila* was digested with KpnI and NdeI and ligated into MCS2 of all the cargo-only expression plasmids, resulting in pETDuet-1/*hpl*/*hrpB_X.nem_* (HGB2529), pETDuet-1/*crpC*/*hrpB_X.nem_* (HGB2528), and pETDuet- 1/*hphA*/*hrpB_X.nem_* (HGB2527). Integration of each T11SS protein was confirmed via digestion with KpnI and NdeI as well as Sanger sequencing using primer 13 the at the University of Tennessee (UT) Genomics Core.

To construct chimeric hemophilin homologs, *hrpC* and *hpl* were split into two domains based on multiple sequence alignment and the NCBI conserved domain database. *hrpC* was split between position 402 and 403, while *hpl* was split between position 474 and 475. Primers 14-15 were used to amplify the hemophilin handle domain from pETDuet-1/*hrpC*/*hrpB_X.nem_* (HGB2530). Primers 16-17 were used to amplify *hrpB* and the hemophilin β-barrel domain from pETDuet-1/*hrpB_X.nem_*/*hpl* (HGB2529). These two products were assembled into pETDuet- 1/Chimeric hemophilin(*hrpC*-*hpl*)/*hrpB_X.nem_* (HGB2595). Primers 18-19 were used to amplify the hemophilin handle domain from pETDuet-1/*hrpB_X.nem_*/*hpl* (HGB2529). To amplify *hrpB_X.nem_* and the hemophilin β-barrel domain from pETDuet-1/*hrpC*/*hrpB_X.nem_* (HGB2530). These two products were assembled into pETDuet-1/Chimeric hemophilin(*hpl*-*hrpC*)/*hrpB_X.nem_*(HGB2596). Primers 14-15 were used to amplify the hemophilin handle domain from pETDuet-1/*hrpC*/*hrpB_X.nem_* (HGB2530). Primers 16-17 were used to amplify *hrpB_H.haem_* and the hemophilin β-barrel domain from pETDuet-1/*hpl*/*hrpB_H.haem_* (HGB2523). These two products were assembled into pETDuet-1/ Chimeric hemophilin (*hrpC*-*hpl*)/*hrpB_H.haem_* (HGB2597). Finally, pETDuet-1/Chimeric hemophilin (*hpl*-*hrpC*)/*hrpB_X.nem_*(HGB2596) and pETDuet-1/*hpl*/*hrpB_H.haem_* (HGB2523) were digested with NotI and NcoI, liberating the hemophilin homolog from each vector. The chimeric hemophilin from HGB2596 was isolated via gel electrophoresis and then ligated into MCS1 of the vector isolated from HGB2523, resulting in pETDuet-1/Chimeric hemophilin (*hpl*-*hrpC*)/*hrpB_H.haem_* (HGB2598). Integration of each T11SS protein and chimeric cargo was confirmed via Sanger sequencing using primers 12-13 at the University of Tennessee (UT) Genomics Core.

### Protein expression and Immunoblotting

*E. coli* strains used for expression experiments were taken fresh from storage at -80°C for each experiment. Strains were cultured on glucose minimal media plates + ampicillin overnight. For each biological replicate, 10 colonies were pooled and inoculated into 5ml of fresh minimal media glucose + ampicillin broth and incubated rotating overnight. Each replicate of each strain was rinsed 2x in PBS and normalized to an OD_600_ of 0.05 in 60ml of glucose minimal media + 1% LB + ampicillin. These were grown shaking at 225rpm until they reached mid log growth (OD_600_ ≈ 1), typically between 5 and 8.5 hours. Upon reaching midlog growth, 25mls of each culture was removed and used as an uninduced T0 control. The remaining 35mls were supplemented with IPTG to a concentration of 0.5mM. One milliliter supernatant samples were taken at 1 and 2.5 hour(s) post induction. Supernatant samples were clarified via centrifugation and filter sterilized. At 2.5 hours post induction the remaining cultures were concentrated via centrifugation, rinsed 2x in PBS, and lysed via sonication (30 s at ∼500-rms volts). Supernatant samples and cellular lysate samples were supplemented with PMSF (1.7 μg/ml), Leupeptin (4.75 μg/ml), and Pepstatin A (0.69 μg/ml) to inhibit proteinase activity. The no plasmid control was performed identically except without the presence of ampicillin in the media.

The protein concentration of cellular lysates was normalized via the Pierce 660nm Protein Assay (REF22660). For supernatant samples 600μl of each filtered sample was precipitated via 10% Trichloroacetic acid precipitation as previously described [12,53]. Samples were boiled for 10-25 min. prior to performing SDS-PAGE to ensure complete unfolding in the protein sample buffer. SDS-PAGE was performed in duplicate using 10% polyacrylamide gels. The first gel was used to perform Coomassie staining for total protein content while the second gel was transferred to a PVDF membrane for Western immunoblotting. Immunoblots were incubated in 50% Ly-cor blocking buffer:50% Tris buffered saline (TBS) for 1 hour to block. Immunoblots were then incubated in 50% Ly-cor blocking buffer:50% TBS supplemented 0.1% Tween20 and 1:5000 rat α-FLAG antibody for 1 hour. Subsequently the blots were incubated in 50% Ly-cor blocking buffer:50% TBS supplemented 0.1% Tween20 and 1:5000 goat α-rat antibody bound to a 680CW fluorophore for 1 hour. Finally, immunoblots were visualized using a Li-cor odyssey imaging the 700nm wavelength. The intensity of supernatant samples was normalized to a clearly visible, non-target protein band in the Coomassie stain to control for protein concentration. Efficacy of secretion was measured as the fold change of cargo protein present in the supernatant when co-expressed with a T11SS protein relative to cargo protein present in the supernatant when expressed alone. Fold changes were not normally distributed initially, so they were log_10_ transformed prior to analysis. The resulting data were analyzed via a one-way ANOVA and a Tukey’s HSD Test [54].

### Purification of hemophilin homologs

Hemophilin from *H. haemolyticus* was expressed and purified as previously described to yield low and high heme-content fractions after anion exchange chromatography [18]. Heme was removed by cold acid acetone treatment to yield an apo hemophilin fraction, as previously described [55]. Residual Fe(III) heme was estimated at 1.8% of sites, based on extinction coefficients of met-hemophilin being 96,100 M^−1^ cm^−1^ and 38,600 M^−1^ cm^−1^ at 414 nm and 280 nm, respectively, and extinction coefficient of the apo-protein being 25,900 M^−1^ cm^−1^ at 280 nm.

Expression constructs encoding the hemophilin homologs from *X. nematophila* (residues 23–247), *X. cabanillasii* (residues 23–238) and *A. baumannii* (residues 21–264) were constructed in pET28a. In each case, the native N-terminal signal peptide was omitted and replaced with a hexa-histidine tag and engineered tobacco etch virus (TEV) protease cleavage site. Clones were transformed into *E. coli* strain Rossetta-2 (Novagen), grown in LB containing 34 μg/ml chloramphenicol and 25 μg/ml kanamycin; expression was induced with 1 mM IPTG for 3 hours shaking at 37°C. Cells were suspended in lysis buffer (0.5 M NaCl, 0.05 M sodium phosphate, 0.02 M imidazole, 100 μM phenylmethylsulfonyl fluoride, pH 7.2) and lysed by sonication (Branson), then hemophilin homologs were captured by Ni-affinity chromatography. TEV protease was expressed and purified as described [56]. The His-tag was cleaved from hemophilin homologs by TEV protease treatment overnight at room temperature, to liberate hemophilin proteins with an additional N-terminal Gly-His-Met tripeptide residual from the TEV cleavage site. TEV protease and His-tag peptides were removed over a second Ni-affinity column. Hemophilin preparations from *X. nematophila* and *A. baumannii* had a brownish appearance and an absorbance peak at ∼413 nm characteristic of a porphyrin ligand, as well as less intense absorption peaks at 533 and 659 nm. Ligand was estimated to occupy ∼25% of sites based on comparison with spectra of hemophilin from *H. haemolyticus*. In contrast, CrpC from *X. cabanillasii* was colorless. Acid acetone or methyl ethyl ketone extraction was not effective to remove colored contaminants from HrpC of *X. nematophila* or *A. baumannii*. Apo-protein fractions of these proteins were prepared by reversed-phase HPLC over a C4 stationary phase (Waters Symmetry) developed with a CH_3_CN:water mobile phase gradient containing 0.1% trifluoroacetic acid. Solvent was removed by lyophilisation. Apo-CrpC from *X. cabanillasii* was applied to a strong anion exchange resin (Q sepharose, Pharmacia) in 25 mM Tris-HCl buffer (pH 8.25 at 21°C) and collected in the flow-through. All apo-proteins were dialyzed into 20 mM Tris-HCl buffer (pH 7.9 at 21°C) prior to storage at –80°C. Apo-protein concentrations were determined by absorption extinction coefficient at 280 nm calculated from amino acid composition.

### UV-visible absorption and fluorescence spectroscopy

UV-visible spectra were recorded on a Jasco V-630 spectrophotometer fitted with a temperature-controlled sample holder (Jasco) and spectrosil quartz cuvettes with a path length of 1.0 cm (Starna, Baulkham Hills, Australia). Porphyrin concentrations were determined according to the following molar extinction coefficients and solvent conditions: hemin chloride, ε_385_ = 58,400 M^−1^cm^−1^ in 0.1 M NaOH [57]; PPIX ε_554_ = 13,500 M^−1^cm^−1^ in 2.7 M HCl [57]; coproporphyrin III, ε_548_ = 16,800 M^−1^cm^−1^ in 0.1 M HCl [58]; Co(III)PPIX ε424 = 180,000 M^−1^cm^−1^ in NaOH(0.1 M):pyridine:H_2_O 3:10:17 [59]. Zn(II)PPIX was determined as free PPIX after decomposition in 2.7 M HCl. To prepare Zn-PPIX, 0.5 g PPIX (Frontier Scientific) was dissolved in boiling chloroform (100 mL), to which a saturated solution of Zn acetate in MeOH (1 mL) was added. The mixture was refluxed for 20 min and then a small amount of MeOH was added and, after cooling, the dark red solid was filtered off (50:50 Zn-PPIX:PPIX by HPLC). Zn-PPIX was purified by RP-HPLC over a C18 solid phase (Phenomenex) with isocratic acetone:MeOH:water:formic acid (280:120:100:1) mobile phase, which achieved baseline separation of the Zn-PPIX fraction.

Hemin binding measurements in absorbance mode were made by successive additions of apo-protein (∼0.4 mM stock) into porphyrin solution (1.5 µM) in 20 mM Tris-HCl, pH 8 at 21°C. All binding experiments were performed at 21°C. Data were fitted to a 1:1 binding model accounting for ligand depletion, where F_obs_ is fluorescence signal, F_0_ is the starting

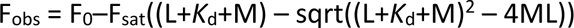

fluorescence, F_sat_ is a scaling factor for fluorescence at saturation, L and M are the ligand and macromolecule concentrations, respectively, and *K*_d_ is the equilibrium dissociation constant. Data were fitted using GNUPLOT version 4.6. A 95% confidence interval for the *K*_d_ parameter was obtained by determining a threshold sum-of-squares for which a fit with all fixed parameters

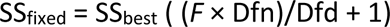

would not be significantly different from the best-fit model at a significance level of P = 0.05 [60]. Here, SS_best_ is the sum-of-squares for the fit with all parameters floated; Dfn and Dfd are the degrees of freedom in the numerator and denominator, respectively, for calculation of *F*. The *K*_d_ value was then fixed at values above or below the best-fit *K*_d_ until fits exceeded the threshold sum-of-squares. To determine a lower limit of Kd that we could expect to fit from absorbance measurements we simulated data for different *K*_d_ values with Gaussian noise added to give fits with SS_best_ that matched our experimental data. We determined by F test that *K*_d_ values ≤ 15 nM were not significantly different (at P = 0.05) from arbitrarily high *K*_d_ (approximating a straight-line fit), thus we decided on 15 nM as a lower cutoff for reporting *K*_d_.

Fluorescence measurements for Zn(II)PPIX binding were made in 96-well format in 20 mM Tris-HCl, pH 8.0 at 21°C using individually mixed samples covering an appropriate range of hemophilin protein concentrations based on preliminary experiments. IC_50_ for hemophilins binding to Zn(II)PPIX (0.5 or 1.0 µM) in competition with BSA (15 µM) were determined by fitted to the same 1:1 binding model as described above. The Cheng and Prusoff equation [61] was used to convert to *K*_d_, where [BSA] is the molar concentration of competing BSA (15 µM)

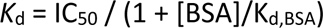

and *K*_d,BSA_ was the dissociation equilibrium constant for BSA binding to Zn(II)PPIX, determined to be 0.3 (0.2–0.4) µM from absorbance titration data.

### Generation of threading models for hemophilin active sites

Threading models were generated with the program MODELLER version 10.2 [32], using the X-ray crystal structures of *H. Haemolyticus* Hpl (PDB 6om5) and *A. baumannii* HphA (PDB 7red) as templates [13,18]. Residues of the N-terminal signal peptide, as identified by SIGNALP 5.0 [62], were removed prior to analysis or modeling of all sequences. The threading approach used by MODELLER relies on accurate alignment of template and query sequences. To identify residues likely to be important for structural integrity of the hemophilin ligand binding domain we collected the small number of sequences that shared similarity with Hpl, HphA and *X. nematophila* HrpC in the N-terminal region, or with pair-wise combinations of these, based on BLASTP searches. We aligned these sequences with the structure-based alignment of Hpl and HphA produced by MODELLER to generate a profile against which subclusters from the sequence similarity network analysis were then aligned using CLUSTALO. Alignments were inspected against several criteria to see that they were structurally plausible. We specifically examined the preservation of the conserved hydrophobic core ligand binding domains of Hpl and HphA, corresponding to Hpl residues G26, I38, I40, A49, V51, I53, F76, M79, A83, A104, L125, F127, Y136, G138, W140, Y136, A158 (Fig. 9). An initial structure-based alignment of *H. haemolyticus* Hpl and *A. baumannii* HphA was produced using MODELLER, and this pairing was fixed in all subsequent alignments. Amino acid residues with potential importance for the hemophilin fold were identified by sequence homology (with the signal peptides and BBD removed) using BLASTP [63]. Sequences were identified with similarity to the hemophilin domains of Hpl, HphA and the N-terminal region of *X. nematophila* HrpC (WP_019473020, WP_057440571, WP_244182492), or Hpl and HphA only (WP_005758278), or HphA and HrpC only (MBP6115507, WP_228864429, WP_232888613), or Hpl and HrpC only (WP_038256617, WP_092512525, WP_244182492), or HphA only (WP_121975315), or Hpl only (RKV63521). These sequences were aligned with the structure-based alignment of Hpl and HphA using CLUSTAL OMEGA [64] to generate a. A profile was generated from this sequence alignment and this profile was aligned with the profiles of the hemophilin network clusters, Plant/Environmental, HphA-like, HrpC-like and CrpC-like, using CLUSTAL OMEGA. Alignments were colored using MVIEW [65].

## Acknowledgements

We thank Dr. Brianna Atto for useful discussions and Sarah Kauffman for technical assistance.

---

**Supplementary Figure S1.**
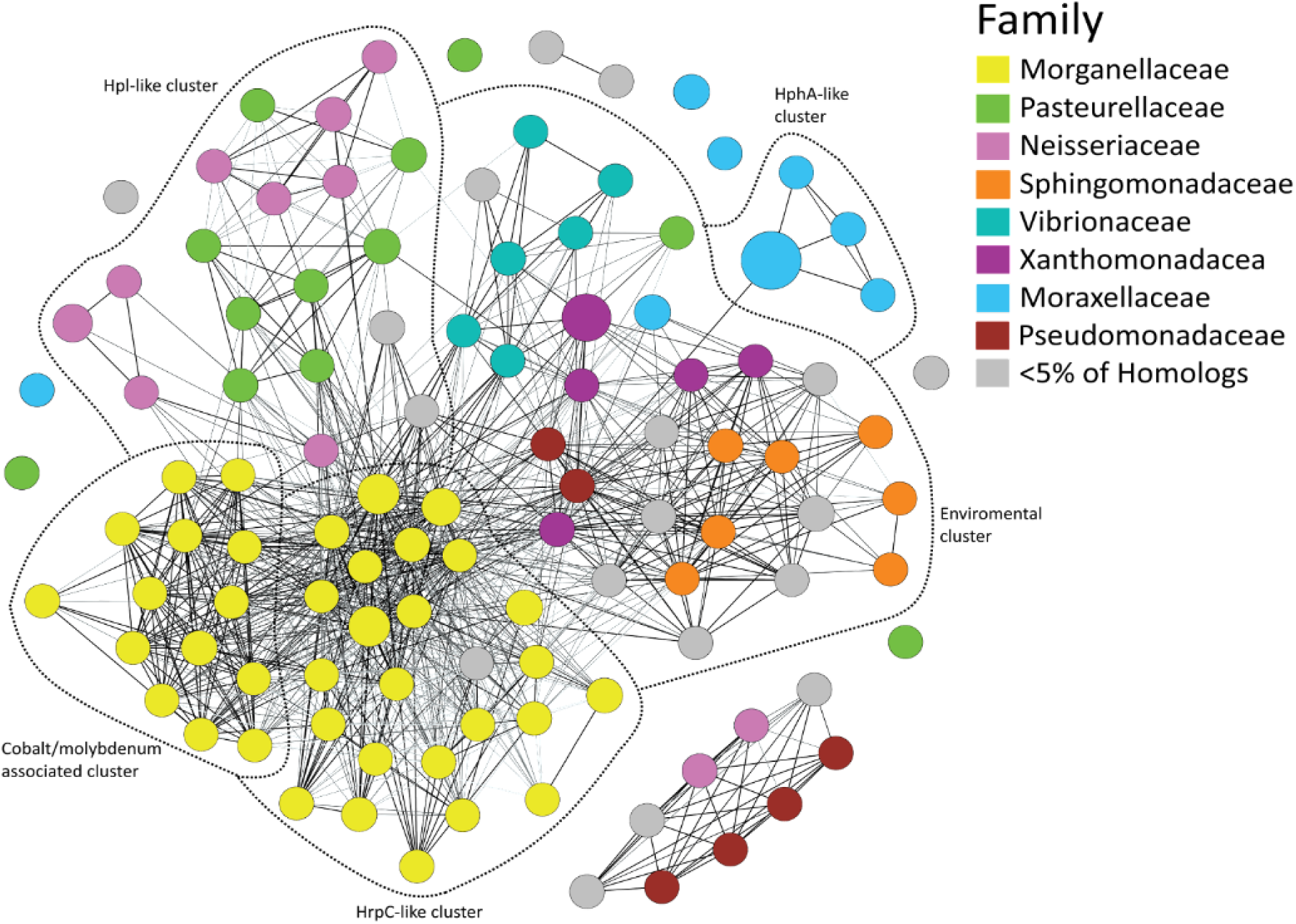
Distribution and relatedness of soluble T11SS-dependent cargo proteins. A sequence similarity network of all soluble TbpBBD-domain-containing proteins identified in Grossman and Mauer, et al. (2022). All of these sequences are encoded in the same genomic region as a T11SS-encoding gene (± 6 genes). Sequences were submitted to EFI-EST (see methods) to generate the network. Each node represents one or more protein sequences with 80% or greater identity, the larger the node the more sequences it contains. Edges indicate an alignment score of 35 or greater. Edge darkness indicates shared sequence identity, with the darkest edges being the most identical. Dotted lines indicate proposed subclusters as defined using the Fruchterman-Reingold algorithm and the distribution of characterized proteins. Sequences that did not connect to any characterized hemophilin family protein were excluded from further analysis (e.g., Figure 3). Despite having similar domain architectures to hemophilin, none of these divergent sequences contained experimentally characterized proteins.

**Supplementary Figure 2.**
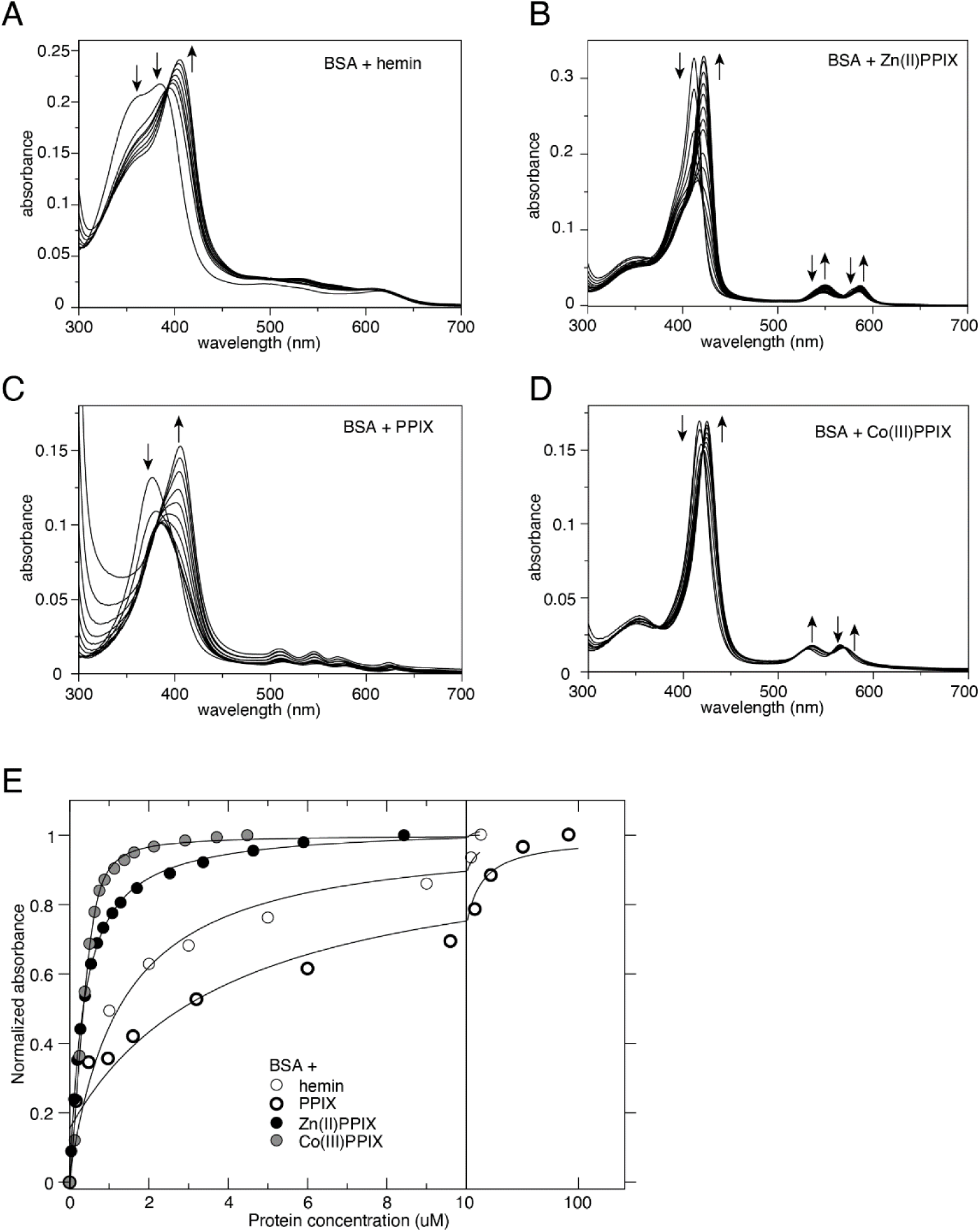
Titration of BSA into porphyrins monitored by UV-vis spectroscopy. **A)** Spectra of hemin (2.5 µM) alone or with addition of BSA to a final concentration covering the range 1–21 µM. **B)** Spectra of Zn(II)PPIX (1.5 µM), alone or with addition of BSA to a final concentration in the range 0.05–12 µM. **C)** Spectra of PPIX (1.5 µM) alone or with addition of BSA to a final concentration in the range 0.2–92 µM. **D)** Spectra of Co(III)PPIX (1.5 µM) alone or with addition of BSA to a final concentration in the range 0.13–4.5 µM BSA. Arrows show the direction of spectral changes. All spectra were recorded in 20 mM Tris-HCl, pH 8.0 (at 21°C). **E)** Binding isotherms for BSA with hemin, PPIX, Zn(II)PPIX and Co(III)PPIX generated from the data in **A–D** by fitting to peak wavelength changes using a 1:1 binding model accounting for ligand depletion.

**Supplementary Figure 3.**
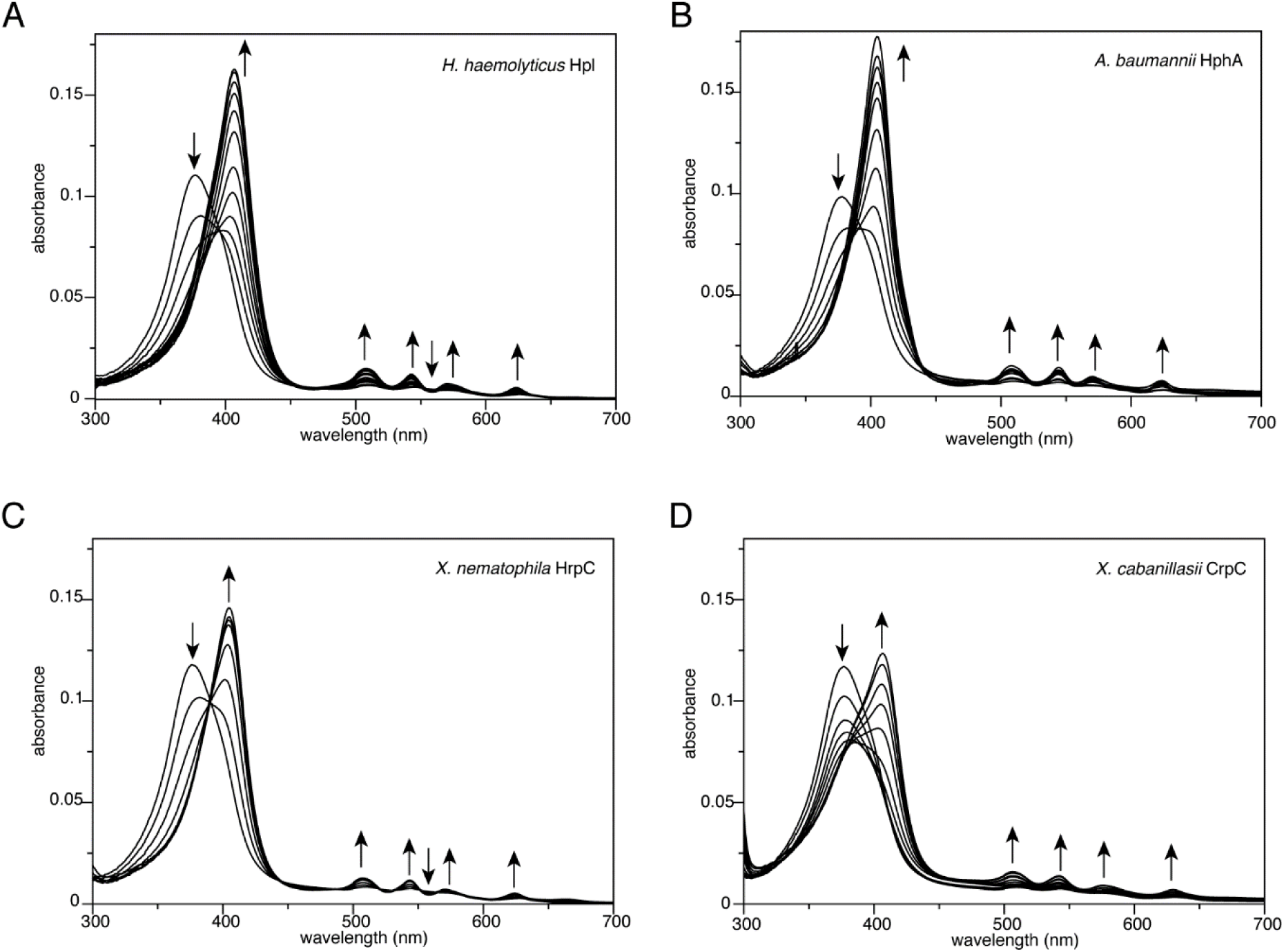
Titration of HrpC proteins in PPIX monitored by UV-vis spectroscopy. **A)** Spectra of PPIX (1.5 µM) alone, or with addition of *H. haemolyticus* Hpl to a final concentration in the range 0.2–4.3 µM. **B)** Spectra of PPIX (1.5 µM) alone, or with addition of *A. baumannii* HphA to a final concentration in the range 0.2–3.3 µM. **C)** Spectra of PPIX (1.5 µM) alone, or with addition of *X. nematophila* HrpC to a final concentration in the range 0.3–3.2 µM. **D)** Spectra of PPIX (1.5 µM) alone, or with addition of *X. cabanillasii* CrpC to a final concentration in the range 0.3–24 µM. Arrows show the direction of spectral changes. All spectra were recorded in 20 mM Tris-HCl, pH 8.0 (at 21°C).

**Supplementary Figure 4.**
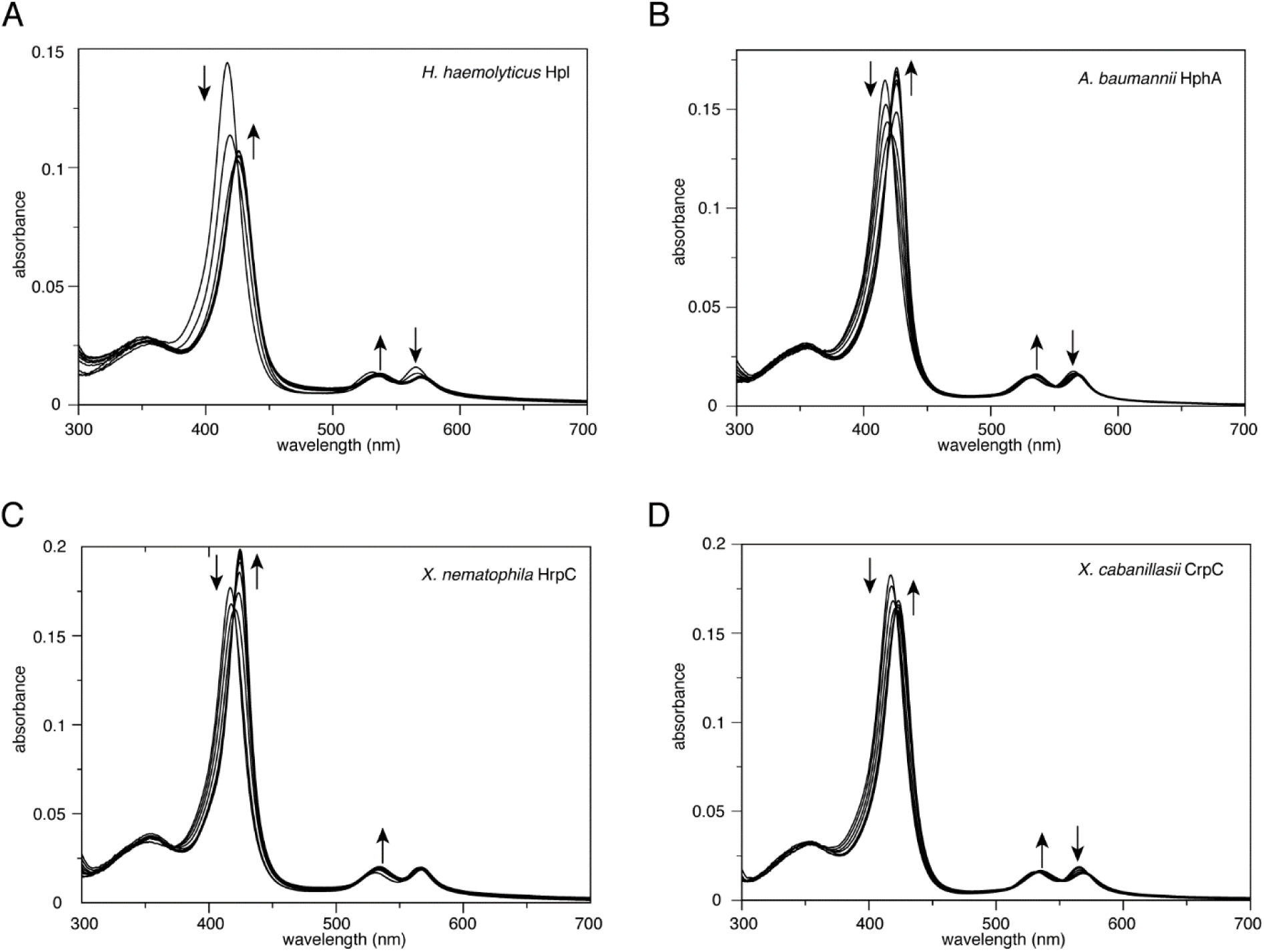
Titration of HrpC proteins in Co(III)PPIX monitored by UV-vis spectroscopy. **A)** Spectra of Co(III)PPIX (1.5 µM) alone, or with addition of *H. haemolyticus* Hpl to a final concentration in the range 0.2–2.2 µM. **B)** Spectra of Co(III)PPIX (1.5 µM) alone, or with addition of *A. baumannii* HphA to a final concentration in the range 0.13–2.6 µM. **C)** Spectra of Co(III)PPIX (1.5 µM) alone, or with addition of *X. nematophila* HrpC to a final concentration in the range 0.3–3.2 µM. **D)** Spectra of Co(III)PPIX (1.5 µM) alone, or with addition of *X. cabanillasii* CrpC to a final concentration in the range 0.15–4.8 µM. Arrows show the direction of spectral changes. All spectra were recorded in 20 mM Tris-HCl, pH 8.0 (at 21°C).

**Supplementary Figure 5.**
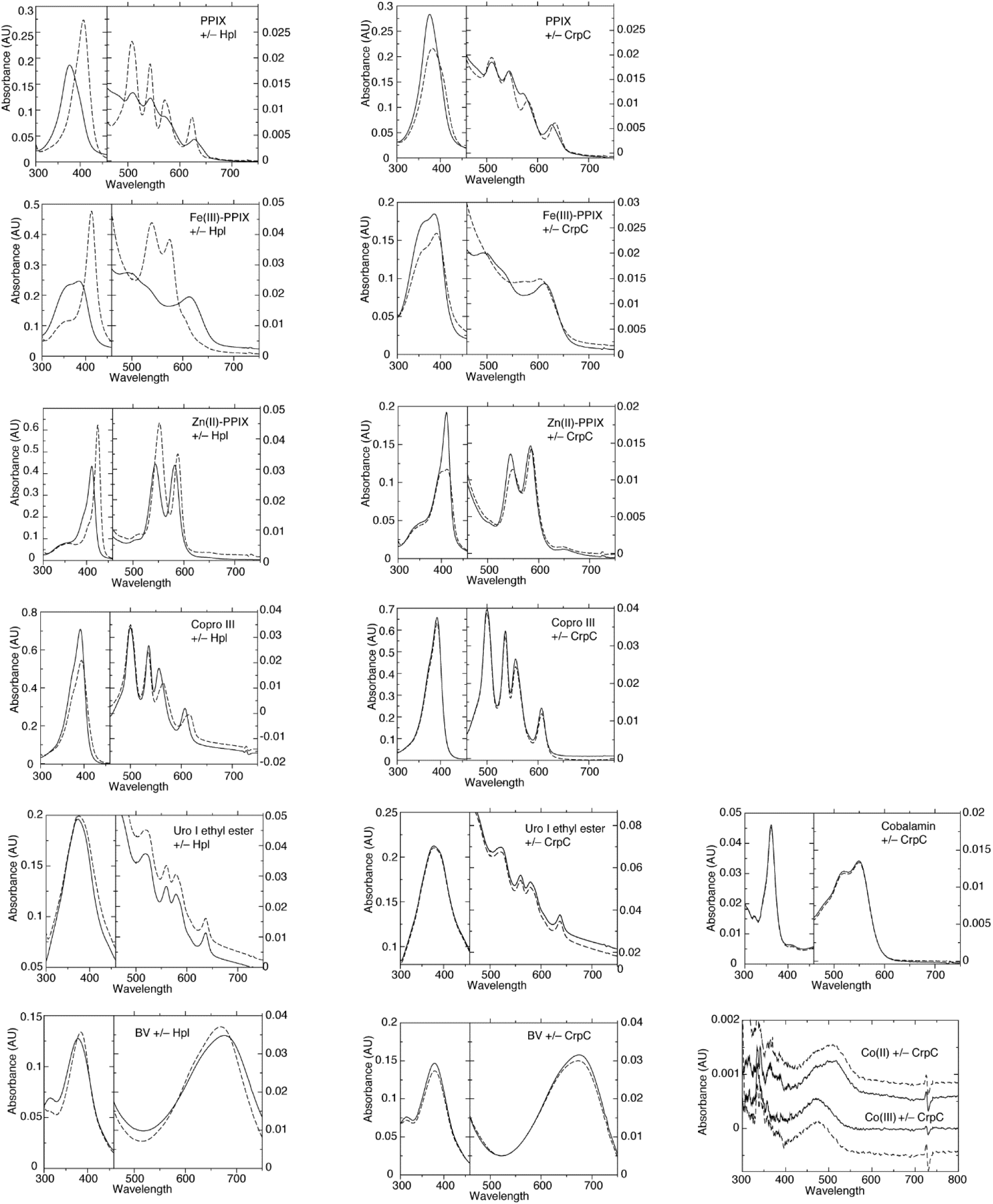
Propensity of Hpl and CrpC to bind to selected linear and cyclic tetrapyrroles. UV-visible spectra of selected tetrapyrroles (4 µM) in the absence (solid lines) or presence of Hpl (left panels, broken lines) or CrpC (right panels, broken lines) at a concentration of 6 µM. Selected ligands are: PPIX, hemin (Fe(III)PPIX), Zn(II)PPIX, Coproporphyrin III (an intermediate in the coproporphyrin pathway of heme synthesis), uroporphyrin I ethyl ester (from porphyric bovine urine), BV (a linear tetrapyrrole and a natural breakdown product of hemin), cobalamin, Co(II) chloride, and hexamine Co(III) chloride. The buffer condition was 20 mM Tris-HCl, pH 8.0 (21°C). Addition of Hpl to PPIX, Zn(II)PPIX, Fe(III)PPIX caused large spectral peak shifts consistent with stoichiometric binding. By comparison, minor changes in absorption peak wavelengths accompanied addition of CrpC to PPIX, Zn(II)PPIX and Fe(III)PPIX reflecting the weak binding characterized in titration studies. Copro III contains four propionate groups and is therefore larger and more polar than PPIX. Weak binding of Hpl to copro III was indicated by small spectroscopic shifts, whereas no binding was evident with CrpC. Similarly, small spectral changes indicated some propensity for Hpl, but not CrpC, to bind to the linear tetrapyrrole, BV. Neither Hpl nor CrpC showed any indication of binding to uroporphyrin I ethyl ester (compared to PPIX, this is a larger amphipathic porphyrin with four alkylated propionates). No spectroscopic changes were induced upon mixing CrpC with cobalamin, Co(II) chloride, and hexamine Co(III) chloride.

**Supplementary Figure S6.**
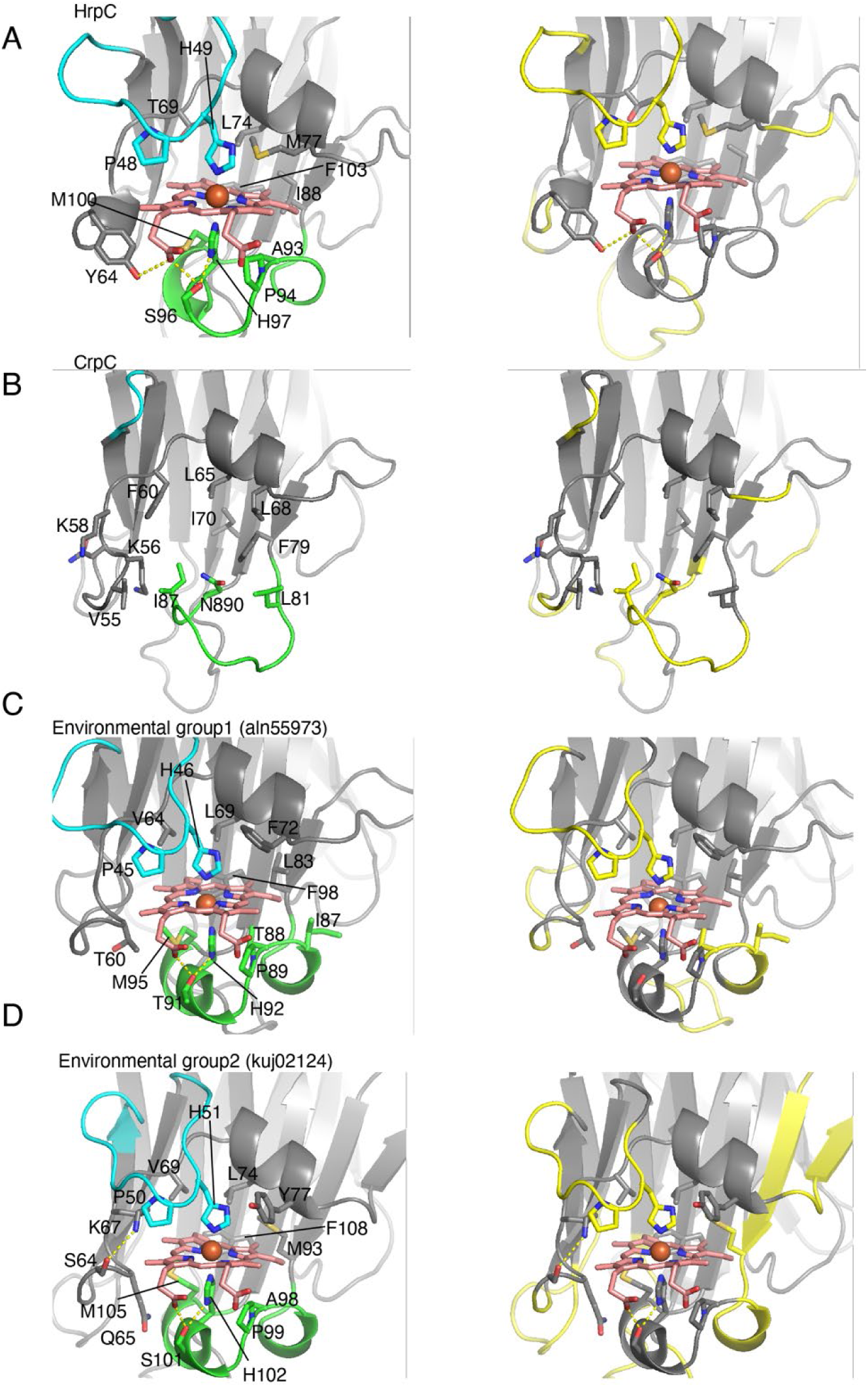
Structural models of hemophilin homologues. Structural model of the hemophilin ligand binding domain are shown for sequences in Figure 9. In the left-hand panels, residues of the β2-β3 (cyan) and β4-β5 (green) loop are shown as in Figure 8. In the right-hand panels, regions of the structure/sequence that do not have sequence-similar threading templates from either *H. haemolyticus* Hpl or *A. baumannii* HphA are colored yellow.

Supplemental file 1: **SubclusterRodeoResultsAndNetworkNodeTable.xlsx** Node information from a sequence similarity network of predicted unlipidated T11SS-dependent cargo proteins generated using EFI-EST, alongside the results of a co-occurrence analysis of hemophilin family subclusters generated using RODEO.

Supplemental file 2: **WesternImages.xlsx** Images of immunoblots performed to demonstrate secretion of hemophilin homologs and chimeric hemophilin acquired using a Li-cor odyssey at 700nm.

Supplemental file 3: **StrainAndPrimerTable.xlsx** Collated descriptions of the bacterial strains utilized within this study and the oligonucleotides used for cloning and sequencing.

